# Benchmarking Framework to Catalyze Individual Human Genome Projects

**DOI:** 10.1101/2025.11.12.688148

**Authors:** Manjushri Kalpande, Apoorva Ganesh, Subhashini Srinivasan

## Abstract

Individual human genome projects still aim for chromosome-level gapless assemblies, which rely on high-coverage reads from multiple long-read sequencing platforms using a multiplicity of assembly pipelines. Moreover, the dependence on DNA derived from primary cell lines for these assemblies makes such projects prohibitively expensive to scale for individual genome initiatives and to catalyze clinical applications.

Over the past decades, genome assembly quality has advanced remarkably from draft assemblies in the early 2010s, to chromosome-level assemblies using error-prone long reads in the late 2010s, to the recent T2T gapless assemblies enabled by high-quality next-generation long-read technologies. That said, a systematic evaluation of trade-offs from assemblies obtained at various coverages, starting at 3x, from a single long-read sequencing platform, is critical for developing a cost-effective and practical strategy for catalyzing individual genome initiative.

Here, by assembling contigs at various coverage levels through downsampling of existing PacBio HiFi reads from three individuals, we demonstrate that high-quality assemblies, as measured by standard assembly metrics and DNA-level linearity relative to a reference across most chromosomes (developed inhouse), can be achieved at approximately 12x coverage. Interestingly, starting at coverages as low as 6x, assembly metrics, including BUSCO scores and DNA-level linearity, begin to saturate, suggesting minimal trade-offs. Furthermore, we show that known structural variants (e.g., the 8p23.1 inversion) can be reliably identified even at 6x coverage.

Together, these results suggest that cost-effective strategies can be developed to advance individual genome initiatives potentially from PacBio HiFi reads from a single SMRT cell per human genome.

## INTRODUCTION

Long-read single-molecule sequencing technologies have not only enabled chromosome-level assemblies for complex genomes but have offered significant reduction in the cost and time. Yet, to catalyze personal genome and biodiversity projects, the cost has to be drastically reduced. Recent studies have addressed the persistent challenges posed by high cost of sequencing, computational demands from high coverage, and use of non-standard analysis pipelines in genome assemblies. Several reports in the recent past have attempted to assess assembly quality using minimal coverage for various long-read technologies using diverse pipelines. Zhang et al assembled *Saccharomyces cerevisiae* and assessed the two long-read sequencing technologies, such as PacBio HiFi and ONT using various assembly strategies, demonstrating the advantages of long-reads for generating accurate contig-level assemblies [1].

Cosma et al systematically compared long-read assemblers across diverse eukaryotic genomes including *S. cerevisiae* (strain S288C), *P. falciparum* (isolate 3D7), *C. elegans* (strain VC2010), *A. thaliana* (ecotype Col-0), *D. melanogaster* (strain ISO-1), and *T. rubripes* and highlighted the importance of assembler choice and read length, showing that longer reads often improve contiguity but not always overall correctness without addressing coverage [2]. According to this report, Flye was the top performer for PacBio CLR and Oxford Nanopore (ONT) reads, while Hifiasm was optimal for PacBio HiFi reads, consistently delivering the best results in terms of assembly quality and completeness across both real and simulated datasets.

Other efforts have focused on evaluating various assemblers specifically on PacBio HiFi sequencing data [3]. In this report the authors have evaluated the performance of 11 de novo HiFi assemblers across multiple plant and animal genomes of varying sizes and complexities. Their results demonstrated that PacBio HiFi sequencing enables near-finished-quality assemblies and identified Hifiasm as one of the topranking assemblers. Yet, like many benchmarking studies, this evaluation was performed under relatively high sequencing coverages (around 30×).

Lang et al used rice samples to generate PacBio HiFi and ONT Nanopore reads [4]. And they report that PacBio Hifi reads were effective at minimising single nucleotide and small insertion/deletion errors, while ONT ultra long reads were shown to help resolve long repetitive regions.

Gavrielatos et al has benchmarked various long-read technologies including Illumina-Nanopore hybrid, Nanopore-only, and PacBio HiFi and respective assembly pipelines such as CANU, HiCanu and Hifiasm for obtaining optimal chromosome-level assemblies [5]. The study concludes that PacBio HiFi sequencing enabled development of novel assembly algorithms requiring less coverage. Furthermore, the use of Hi-C data helps produce long scaffolds of contigs from lower coverage. Accordingly, they report that for human genome assemblies of significant quality can be achieved with coverage as little as 16× of PacBio HiFi data along with use of 10× Bionano and Hi-C data for scaffolding in higher and complex organisms.

Wang et al. studied various assembly strategies and sequencing technologies, such as PacBio HiFi versus ONT, to benchmark the best approach for chromosome-level genome assembly from human samples [6]. The study used publicly available datasets (NA12878 and NA24385) from Genome in a Bottle (GIAB) consortium. The authors demonstrated that assemblies generated using PacBio HiFi reads for diploid genomes showed exceptionally high breadth of coverage, superior accuracy and very low error rates. Furthermore, the incorporation of long range DNA interaction data, especially Hi-C sequencing data, was shown to be indispensable for scaffolding and achieving chromosome-scale contiguity.

The Human Pangenome Reference Consortium (HPRC) released a high-quality contig-level assemblies from 47 diploid individuals using long-read sequencing technologies, primarily PacBio HiFi and Hi-C although reads from other platforms were generated [7]. This work demonstrated that contig-level assembly is sufficient to capture hundreds of megabases of previously unrepresented euchromatic sequences and structural variants missing from the GRCh38 reference. Sarashetti et al. evaluates optimal data types and coverage requirements to generate high quality haplotype-resolved genome assemblies suitable for population-level pangenome projects [8]. In this report, the authors have examined the importance of long-reads from various platforms in the context of achieving high-quality phased genomes with enhanced contiguity and completeness. They have also downsampled to show that a minimal coverage of 20× of PacBio HiFi reads along with ultralong reads from ONT and Hi-C data is sufficient to produce haplotype resolved assemblies.

Taken together, the consensus from benchmarking studies reviewed above, suggests that contigs from PacBio HiFi longreads using Hifiasm assembler scaffolded by Hi-C reads is optimal to obtain high-quality chromosome-level assemblies. Since the computational cost in obtaining contig-level assembly raises exponentially because of high RAM requirements resulting from non-paralizable algorithms, in this study, we address the lowest PacBio HiFi coverage needed for scaffolding with Hi-C data to generate chromosome-scale human genome assemblies with acceptable trade-offs.

## RESULTS

To evaluate the minimal coverage requirement for PacBio HiFi reads, originally sequenced at >32× coverage and available in the public repository at IGSR, from three individuals of South Asians descent (PJL1, BEB1, and ITU1), were downsampled. Assemblies were generated at 3×, 6×, 9×, 12×, and 15× coverage to systematically assess the impact of sequencing depth on chromosome level assembly and respective trade-offs.

At 3× coverage of PJL1, contig-level assemblies were highly fragmented (N50 = 0.095MB; L50 = 7340; Table 1), reflecting poor contiguity for scaffolding using Hi-C as reflected in the N50 and L50 at scaffold level (N50 =0.094MB; L50 =7466; Table 1). Consequently, 3× coverage was excluded from further consideration.

**Table 1:**
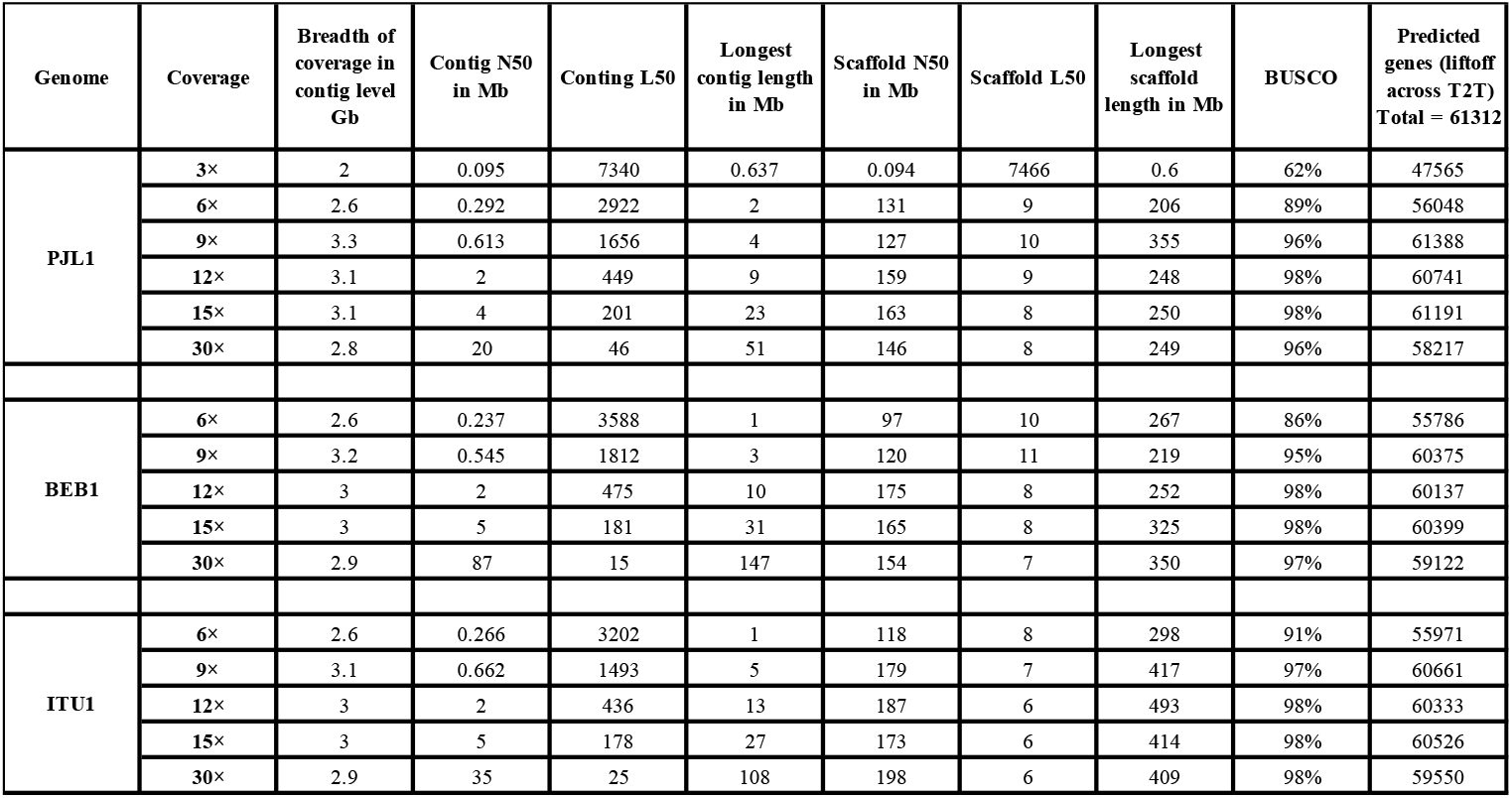
Assembly Statistics Across Genomes at Varying Coverage.

From 6× coverage onwards, contig-level assembly quality based on assembly metrics improved consistently across the genomes of all three individuals. For example, for PJL1, contig-level N50 values rose uniformly from sub-megabase lengths at 6× to tens of megabases at 15×, while L50 values decreased respectively from 2922 to 201. Interestingly, the N50 values for contigs from coverage as low as 6× could be scaffolded to achieve saturation in N50 value after scaffolding using Hi-C. For example, even at 6× coverage the scaffold-level L50 drops to ∼9 with N50 jumping to hundreds of megabases, which is close to the expected N50 of 147 Mb and L50 of 8 achievable for the human genome.

The assembly metrics stated above such as N50, L50, longest scaffolds fail to fully evaluate the quality of assemblies through the length of all 23 chromosomes for complex genomes like that of humans. A DNA-level linearity through the lengths of chromosomes compared to a reference is critical to offer a better measure of quality of assemblies at each coverage. For this, we have used 2kb virtual markers for human derived from two references-hg38 (hg38100kb) and T2T-CHM13v2.0 (T2T100kb) at 100kb intervals and mapped these to assembled scaffolds from each coverage to both assign chromosomes to scaffolds and to establish linearity at the DNA-level [9]. We find that at 12× coverage, most chromosomes were fully collinear with the hg38 reference in all three individuals, except for chromosome 16 in BEB1 and ITU1, and chromosomes 10 and 22 in PJL1, which reflects localized breaks or inversions. However, by 15× coverage, all chromosomes were fully collinear, showing that near-complete assemblies occur by 12× and full structural resolution is obtained by 15×.

Figure 1a summarizes the linearity (dark green), completeness (light green), fragmentation (yellow) and local orientation (pink) in assemblies from three individuals across 4 coverages from a visual assessment of dot plots shown in Supplementary Figure S1:a-w generated using virtual markers through the lengths of the chromosomes in all assemblies compared here. Figure 1c gives examples of four types of scaffolding patterns used to summarize the quality of DNA-level completeness in Figure 1a.

**Fig 1:**
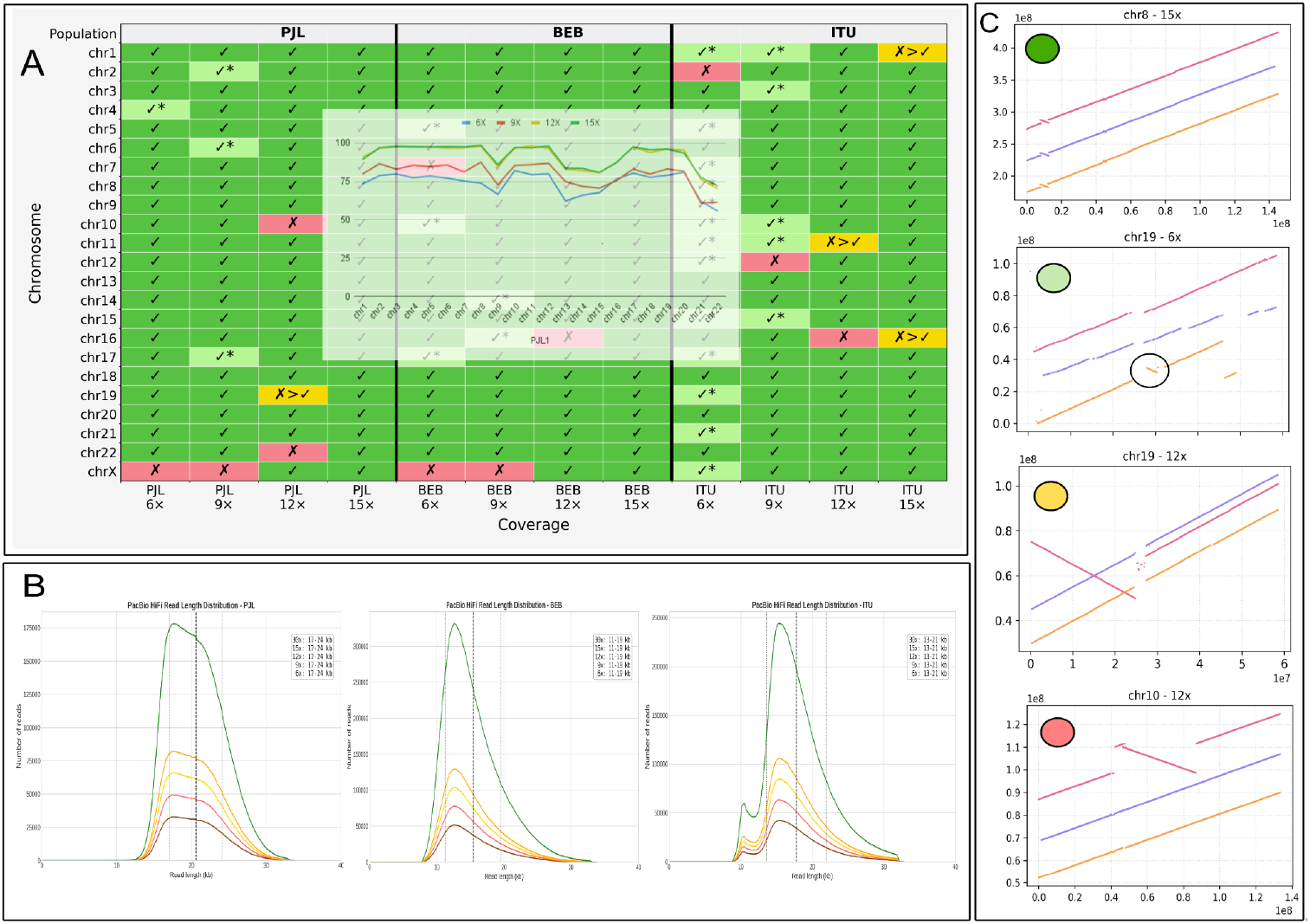
A) Chromosome-wide summary of dot plots shown in Supplementary Figures S1a-w at coverages 6×-15× for individuals including PJL1 (left panel), BEB1 (middle panel) and ITU1 (right panel). Cells in dark green end-to-end linearity with virtual markers, light green are partial linear, yellow manually correctable, and pink with inversions. Inset, percentage of virtual markers from hg38 reference hitting top two scaffolds. B) read length distribution for source data (green) and for all downsampled datasets (6× - brown, 9× - red, 12×-yellow and 15×-orange) for PJL1 (top), BEB1 (middle) and ITU1 (bottom). C) examples of dot plots for different colors shown in A.

To quantitate the DNA-level linearity we computed the percentage of virtual markers hitting the top two scaffolds of each chromosome for every assembly across various coverages for PJL1. In Figure 1a-inset we show the percentage coverage of markers by chromosome for assemblies at different coverages (6×-blue, 9×-red, 12×-yellow, 15×-green). We observe that at 12× a majority of the chromosomes (∼98%) maps to virtual markers with the exception of acrocentric chromosome (chr13, chr14, chr15, chr21, chr22) and chromosome 9 with huge centromere as expected, which is the same at 15×.

We do find significant discrepancy in assemblies at the DNA-level linearity at lower coverage from different individuals. For example, the number of dark versus light green boxes in 6× coverage between PJL1 and ITU1. To explain the source of such discrepancy we checked the PacBio HiFi read length distributions for the three individuals. The read length profile at all coverages were consistent with the original data after downsampling. We find that the read length distribution ranging from 17-24kb in PJL1 may be optimal for assembly at lower coverages. However, the read length distribution in ITU1 not only ranges between 13-21kb but it has an additional peak with many short reads as shown in Figure 1b. We conclude that read length distribution may impact minimal coverage requirement.

Figure 2 shows plots of scaffold coordinates on the y-axis mapping to the virtual marker coordinates for chromosome 8 from T2T (Figure 2a) and hg38 (Figure 2b) for assemblies from all 4 coverages assembled for all three individuals including PJL1 (red), BEB1 (blue) and ITU1 (orange). Since it is reported that hg38 lacks this inversion, the inverted genotypes present in all three individuals when compared with hg38100kb (Figure 2c^4^) are visible in the dot plots and are highlighted in Figure 2b. Thus, presence and absence of known inversions are detectable even at coverages as low as 6×, suggesting that high-quality local assemblies at the contig-level is achievable even at 6× coverages except for ITU1.

**Figure 2:**
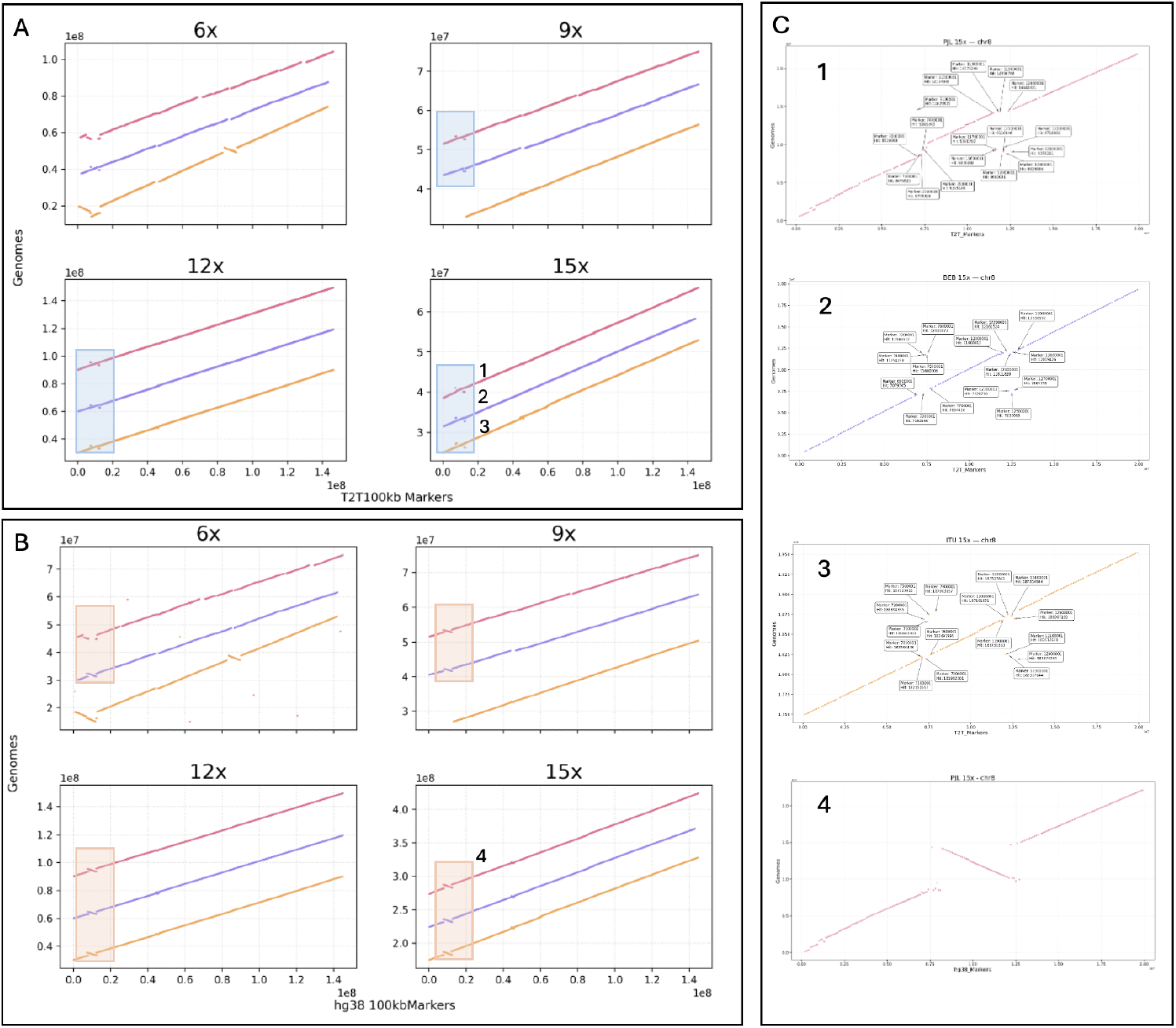
Dot plots showing the linearity of chromosome 8 with respect to: A) T2T100kb and B) hg38100kb virtual markers with panels representing assemblies with coverage ranging from 6× to 15× with individuals PJL1 (pink), BEB1 (purple), and ITU1 (orange). Rectangles in A and B highlight 8p23.1 inversion locus. C) enlarged views of selected loci that are seen off diagonal in 1) PJL1, 2) BEB1, 3) ITU1 from assemblies with 15× coverage and 4) enlarged view of inversion genotype from hg38100kb of PJL1-15×.

**Figure 3:**
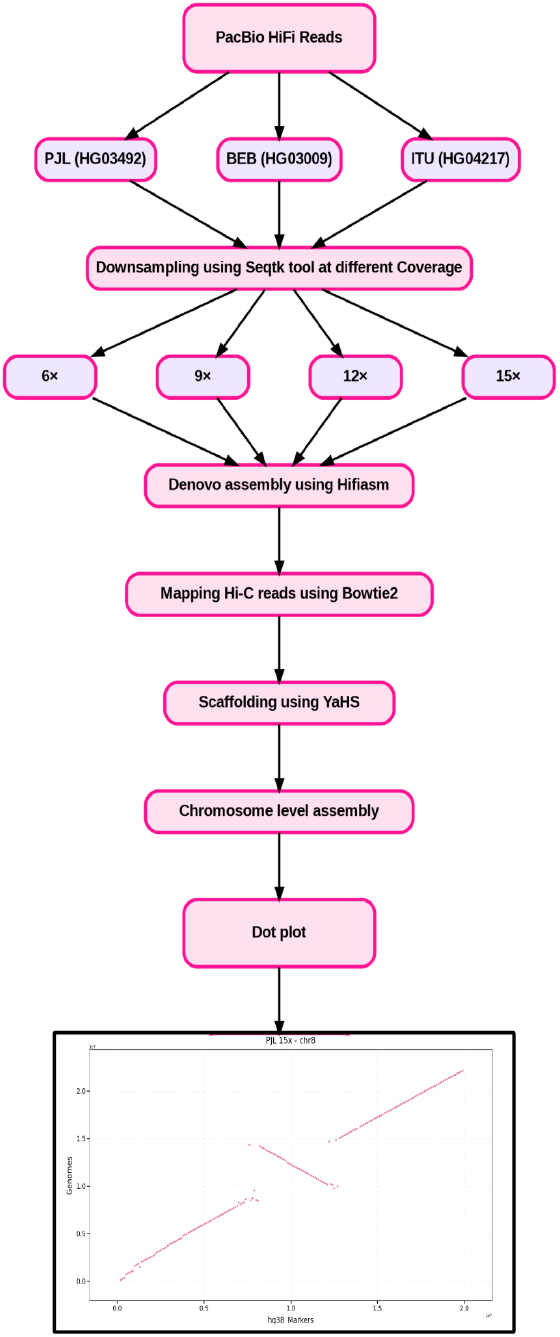
Workflow used for assembly and validation

However, since the inverted genotype between T2T and the three individuals are the same, the dots within the inversion locus align with diagonal as highlighted by rectangles in Figure 2a. Since all three individuals share the same inversion genotype as T2T, we do not expect any off-diagonal markers in this region highlighted in Figure 2a. However, some markers from the 8p23.1 distal inversion breakpoints align with the proximal breakpoint and vice versa, leading to dots that are off-diagonal as shown labeled in Figure 2c^1,2,3^. The variation in the virtual markers at the two 8p23.1 breakpoints among the three individuals suggest 8p23.1 inversion hotspots in humans.

## DISCUSSION

Over the past two decades, genome assembly quality has advanced remarkably starting from i) draft assemblies from Illumina short reads to build contigs and scaffolded by mate-pair reads in the early 2010s, ii) chromosome-level assemblies using Hi-C reads with contigs built using high-coverage error-prone long reads in the late 2010s, and iii) the recent high-quality T2T assemblies enabled by second-generation high-quality (HiFi) long-read technologies. Yet, at every stage, the assemblies were valuable in extracting useful biological inferences. Conventional wisdom suggests that higher the coverage, higher the quality of assembly and higher the cost. There is a need to evaluate minimum coverage of high-quality long-reads at which the assembled genome has least trade-offs. This said, genome assembly benchmarking should include measuring completeness and usefulness, along with standard metrics including N50, L50, longest contig/scaffold size, number of gaps, and synteny preservation. BUSCO score for gene completeness and the preservation of DNA-level linearity of scaffolds with reference chromosomes should be used in benchmarking studies and assessing the trade-offs at each coverage.

As shown in Table 1, by evaluating assemblies at different coverages, we found that at 6× the breadth of coverage by contigs (Table 1, column 3) start approaching the genome size and that the contig-level L50 and N50 are optimal for scaffolding into chromosomes using Hi-C read pairs. However, at 3× coverage, not only is the N50 value small and the L50 large, but scaffolding with Hi-C is untenable, as shown in Table 1. For assemblies with coverage greater than or equal to 6×, contig-level L50 and N50 values improve with increasing coverage, while scaffold-level L50, N50, and BUSCO scores begin to saturate at 6×, suggesting that scaffold-level assemblies from 6× and higher coverage are already useful for many applications.

We developed a virtual marker-based approach to assess the DNA-level linearity of contigs within chromosomes at varying coverages, achieved by scaffolding. From coverages of 6× onwards, the contigs are not only correctly binned into chromosomes by Hi-C reads, but are also stitched linearly for a majority of the chromosomes (Figure 1a), leading to individual scaffolds representing chromosomes. This approach, while assigning scaffolds to chromosomes, also validates the linearity of respective scaffolds at each coverage level with the respective references such as hg38 and T2T. We quantitate DNA-level linearity in Figure 1a-inset by calculating percentage coverage of markers by chromosomes for assemblies at different coverages (6×-blue, 9×-red, 12×-yellow, 15×-green). We observe that at 12× a majority of the chromosomes (∼98%) maps to virtual markers with the exception of acrocentric chromosome (chr13, chr14, chr15, chr21, chr22) and chromosome 9 with huge centromere as expected, which does not improve at higher coverage of 15×. Suggesting that at 12× coverage sufficient DNA-level linearity is also achieved and saturated.

Interestingly, as shown in Figure 1 (dark green), DNA-level linearity and contig orientations for many chromosomes is already validated end-to-end at a coverage of 6× in individuals except ITU1. Major discrepancies are only in assigning correct orientations (Figure 1a, pink) for a few fragments/contigs within its locus at lower coverages, which may be resulting from unresolved strand orientations by Lachesis algorithm while stitching contigs into scaffolds by Hi-C reads when contigs are too short for multiple Hi-C reads to map.

In order to understand the observed disparity in the quality of DNA-level assemblies for some chromosomes at lower coverages between ITU1 and PJL1, we explored the role of read length distribution as shown in Figure 1b. We conclude that the mean read length distribution as in PJL1 may play a significant role in high-level assemblies at lower coverage across chromosomes. Thus, the minimum coverage requirement is dependent on the quality of raw sequencing data, which depends on the experimental condition varying from lab-to-lab. In any event, wrongly oriented DNA fragments/contigs can be identified by mapping PacBio HiFi raw reads onto the reference for any sign of discontinuity of reads mapping to the reference as shown by the author’s lab for unique inversions observed for KIn1 chromosome 7 and 17 [9]. Thus, even assemblies from lower coverages in ITU1 and BEB1 with orientation errors (Figure 1a, pink) can be detected and, if needed, can be manually corrected — a tedious process with only cosmetic benefit. Despite orientation issues, we have shown that large known inversions such as 8p23.1 are detectable at 6× coverage in all three individuals (Figure 2b) by using hg38 as reference. Figure 2a shows 8p23.1 inversion locus using T2T as reference, where only the breakpoints are visibly seen off-diagonal because the genotype of this inversion in all three individuals match with that of T2T. The enlarged views off-diagonal virtual markers are labeled in Figure 2c from assemblies at 15x, which varies between individuals.

The benchmarking framework provided here for the human genome, may need to be re-evaluated for organisms with even more complex genomes. In biodiversity projects one is interested in assembling the genomes of thousands of organisms with no reference. However, the detailed benchmarking provided here for humans could also extend to manage cost in biodiversity and earth genome projects. In biodiversity projects, achieving DNA-level linearity at high costs may not necessarily add more value to a project. The high BUSCO scores at much lower coverages suggest usefulness of assemblies at lower coverages in detecting core versus dispensable genes across species or plant strains. We show that even at 6× coverage, all individuals maintained BUSCO [10] scores of ∼89%, as shown in Table 1. Meaning that much of the gene rich regions are captured in reads and get assembled. The high L50 values at lower coverage may result from repetitive regions. This was true during the era of draft genome assembly when more than 80% of the species specific genes could be characterised [11].

Thus, while there is a temptation to produce gapless genome assemblies as a technical feat without cost consideration, we advocate that at much lower coverages one can produce high-quality useful assemblies at fraction of the cost. Moreover, since 6×-9× coverages for humans can be generated from a single SMRT cell, the cost of individual genome initiative can be further contained with manageable trade-offs.

## MATERIALS and METHOD

### Materials

We downloaded PacBio HiFi reads generated for the *Human Pangenome Reference Consortium (HPRC)* from three individuals, PJL1 (HG03492), BEB1 (HG03009), and ITU1 (HG04217) for a coverage of ∼> 30× each representing distinct South Asian populations. For PJL1, both PacBio HiFi reads and Hi-C data were downloaded from the HPRC archive hosted on Amazon S3, available as ccs.bam and FASTQ files, respectively. For BEB1 and ITU1, raw HiFi reads were downloaded as FASTQ files from the HPRC repository, while Hi-C data were retrieved from the Ensembl-EBI database (Data availability).

### Downsampling for Various Coverages

To estimate total genome coverage, reads from all three SMRT cells were merged using the bcftools merge command-line utility. Coverage was then calculated using the formula Coverage = (read count × read length) / genome size. To evaluate assembly performance across varying sequencing depths, reads were randomly downsampled for N× coverages by computing number of reads required for a given coverage by computing average read length for the original PacBio HiFi read file and using the formula no of reads = N ^*^ genome size/average read length. Downsampling was done using Seqtk (v1.3) with the following command-line –

***seqtk sample -s100 sample*.*ccs*.*fastq*.*gz N > N×_sample_ccs*.*fastq***

Here, -s100 sets the random seed and N is the number of reads to be randomly selected from the original PacBio HiFi file. This random subsampling ensured uniform coverage distribution while maintaining read-length profiles representative of the original dataset (Figure 1b).

### Genome Assembly and Comparative Genome Workflow

Genome assembly was performed following three-steps including contig-level assembly, scaffolding and comparative genomics. De novo contig-level assembly of each downsampled PacBio HiFi reads were done using Hifiasm tool [12] using default parameters. Scaffolding of the resulting contigs were done using Hi-C reads by mapping them on to the assembled contigs using Bowtie2 [13] using default parameter, which was then used in scaffolding using YaHS [14] with default tool parameters into chromosome-level scaffolds.

#### 1. Assessing quality of assemblies at each coverage

We have used standard matrices and DNA-level linearity for each chromosome to select the coverage that worked for the majority of the chromosomes.

#### 2. Computing Assembly Matrices

Across all assemblies both at contig-level and scaffold level, we computed N50, L50, longest contig/scaffold, genome size using inhouse developed script (Data Availability) and BUSCO score using the mammalia_odb10 dataset which contains 9,226 conserved single-copy orthologs. BUSCO analysis was performed using the mammalia_odb10 lineage dataset in offline mode, with miniprot specified as the manually provided aligner for protein mapping using command line -

***busco -i <input_genome*.*fa> -l mammalia_odb10 -o <output_folder> -m genome --cpu 12 - m genome --offline --miniprot***

#### 3. Testing DNA-level linearity in scaffolds

Virtual Markers were generated by selecting 2kb fragment evenly spaced genomic markers every 100 kb along each chromosome of the two reference genomes - hg38 (hg38100kb) and T2T-CHM13v2.0 (T2T100kb) using custom scripts (Data Availability).

To validate and visualize DNA-level linearity in the assembled scaffolds at every coverage, we generated dot plots using inhouse scripts (Data availability) by aligning virtual markers from the hg38100kb and T2T100kb. These were aligned to each assembled genome for all three individuals (PJL1, BEB1, ITU1) at different coverage levels using blastn. Only hits with >99% identity with the virtual markers were retained to ensure exact DNA-level matches, and alignments were further filtered to include only markers mapping to the longest scaffolds. For each retained marker, the start coordinate on the reference (x-axis) and the corresponding start coordinate on the assembly scaffold (y-axis) were extracted to generate dot plots (Supplementary Figure S1 a-w), where dots on the diagonal indicate conserved DNA-level linearity and off-diagonal lines indicate inversions or other assembly artifacts between each assembly and the reference genome.

### Manual Correction

During the chromosome assembly process, several chromosomes initially appeared fragmented due to artificial splits between their *p* and *q* arms. In **chr1_ITU1_15×**, markers from the same chromosome aligned to two separate scaffolds (scaffold 9 and scaffold 2), producing discontinuous regions corresponding to the *p* and *q* arms with respect to the references. Re-blasting these scaffolds and re-evaluating the alignments confirmed that they represented adjacent portions of the same chromosome; merging them restored a single, continuous alignment (Supplementary Figure S1-a). Similarly, in **chr11_ITU1_12×** (Supplementary Figure S1 - k) **and chr16_ITU1_15×** (Supplementary Figure S1 - p), several markers in the intermediate region failed to align because of local discontinuities between the *p* and *q* arms. This was resolved by treating the two arms as independent segments by modifying the code, aligning them separately, and then merging the results using an offset-based approach, which restored continuous marker placement and correct scaffold ordering. Comparable issues were also observed in **chr19_PJL1_12×** (Supplementary Figure S1 - s), where scaffolds 20 and 22 exhibited nearly identical top BLAST hit scores to the same chromosome, creating an apparent split between the arms. Merging these scaffolds and re-running the BLAST alignment restored complete chromosomal continuity across both assemblies.

## ABBREVIATIONS

PJL1: Punjabis from Lahore
BEB1: Bengali from Bangladesh
ITU1: Indian Telugus from UK
HPRC: Human Pangenome Reference Consortium
BUSCO: Benchmarking Universal Single-Copy Orthologs
ONT: Oxford Nanopore Technologies
CLR: Continuous Long Reads

## ACKNOWLEDGEMENT

Data analysis personnel including MK and AG were funded via a BioIT grant from Government of Karnataka and computing infrastructure via the Department of Information Technology, Biotechnology and Science and Technology, GoI. The authors also acknowledge using computing resources obtained as part of DBT Builder Sanction no. BT/INF/22/SP45402/2022 dated March 8, 2022 and CCB Sanction no. BT/PR40212/BTIS/137/40/2022 dated December 19, 2022. The authors would like to acknowledge Prof. Bibha Choudhary for her constant support, scientific insight and managing projects under the BioIT grant.

## DATA AVAILABILITY

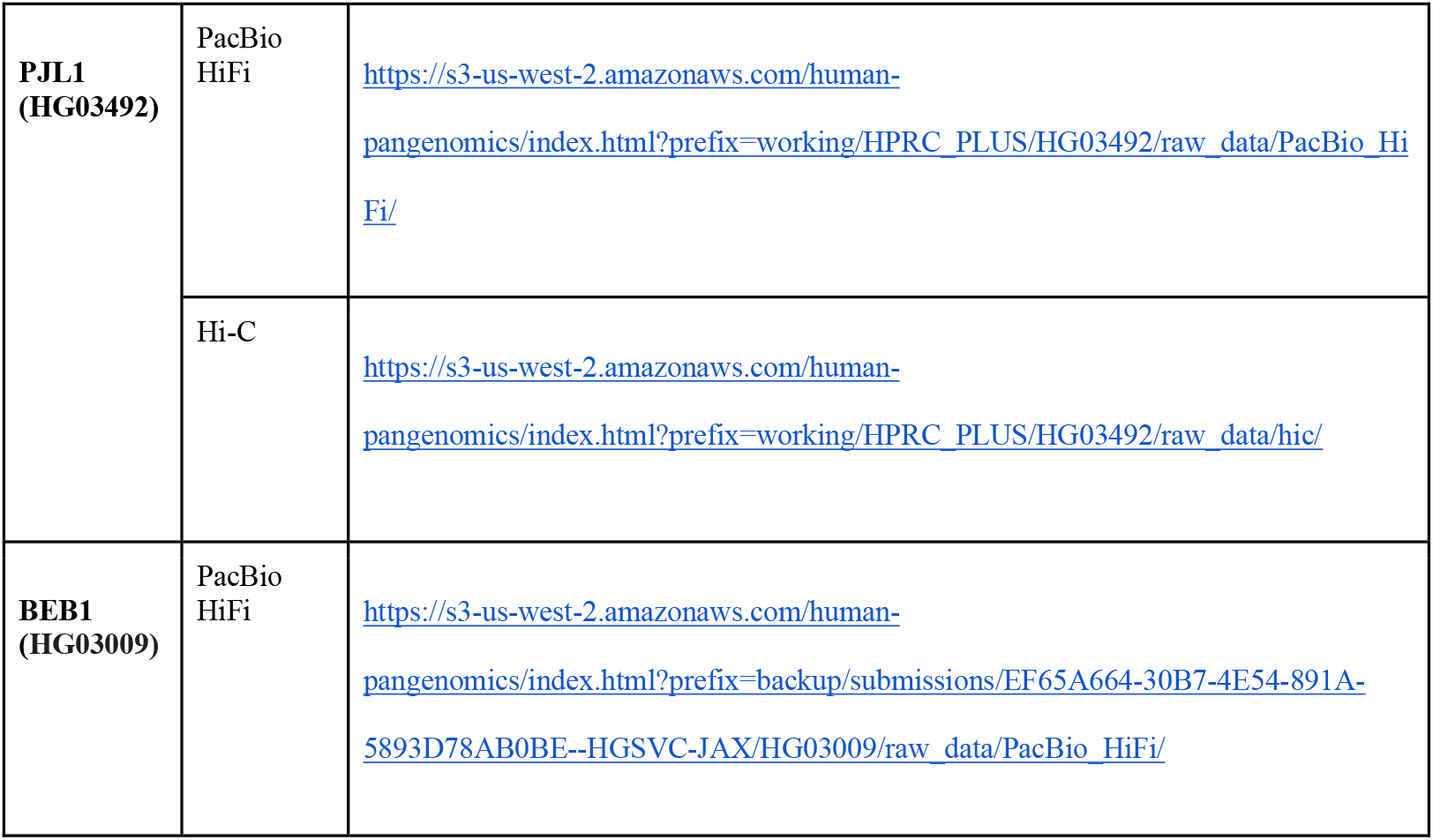
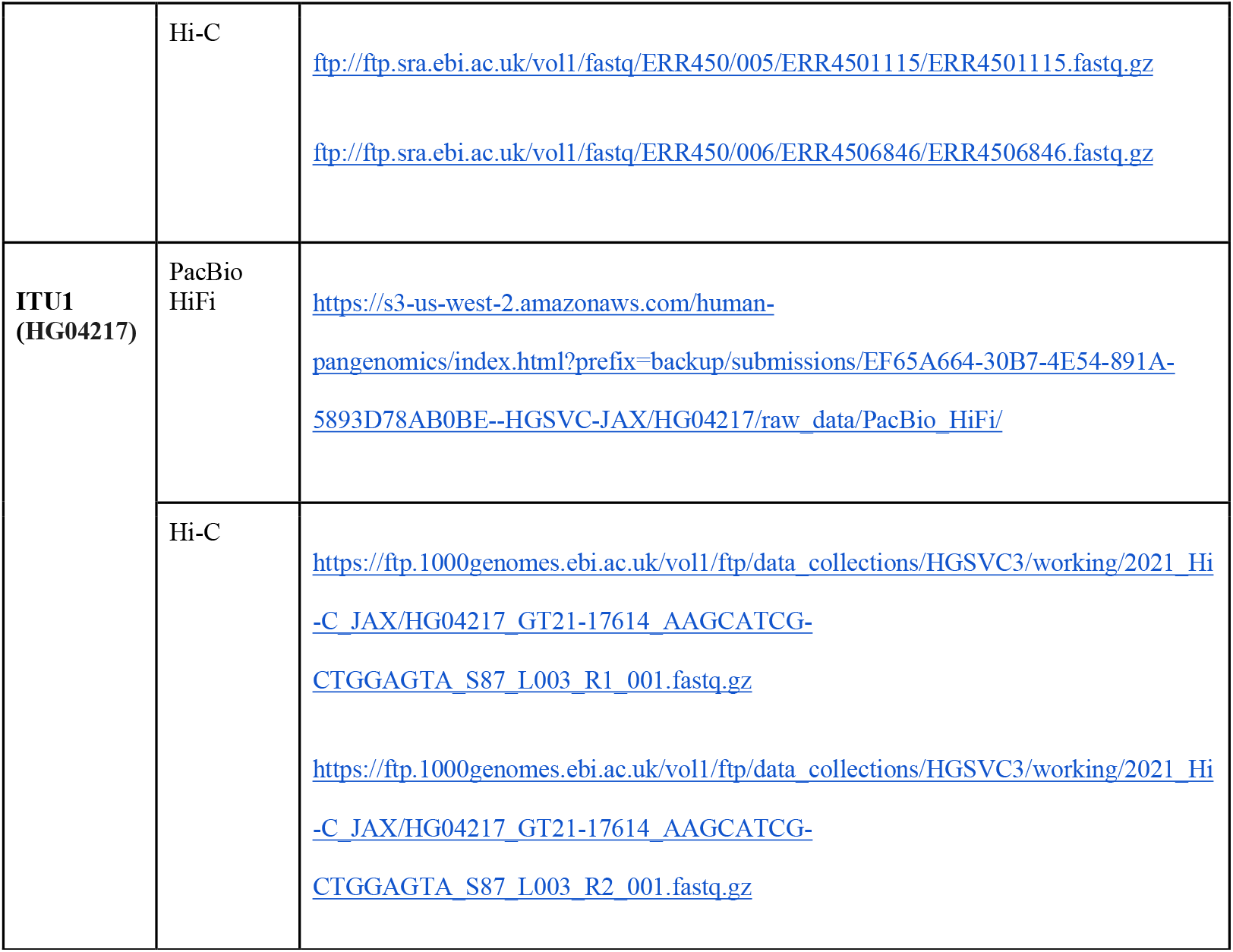

The datasets supporting the conclusions of this article are available in the [**Benchmark_Dataset**] repository, [https://github.com/kalpandemanjushri12/Benchmark_Dataset]

## AUTHOR CONTRIBUTION

MK for generating all the assemblies and generating assembly matrics, AG for writing the manuscript and overseeing the evaluation of scaffold linearity, SS for conceptualizing, designing the experiment and writing of the manuscript

