## Supplementary material for "Benchmarking Framework to Catalyze Individual Human Genome Projects"

Table of Contents

*Supplementary Figure S1 ( A - W )* : shows comparative dot plot of individual chromosomes from genomes ( PJL1, BEB1, ITU1) against virtual markers every 100kb from hg38 assembly……………………………………………………………………………………...	13

*Supplementary Figure S2 ( i - iv )* : shows comparative dot plot of individual chromosomes from few genomes in which p and q arms are splitting ……………………………………14

#chromosome 1

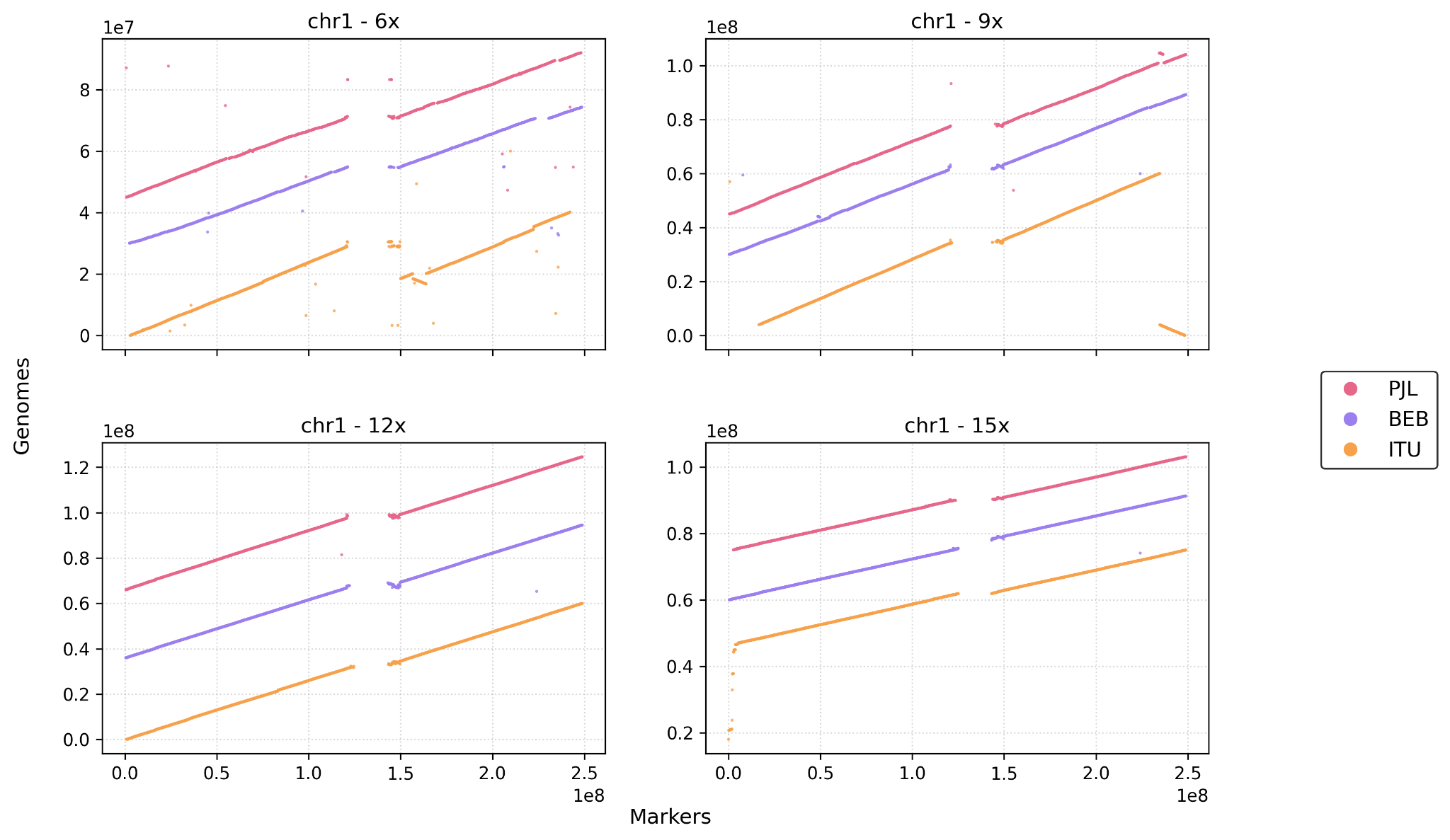

(A)

#chromosome 2

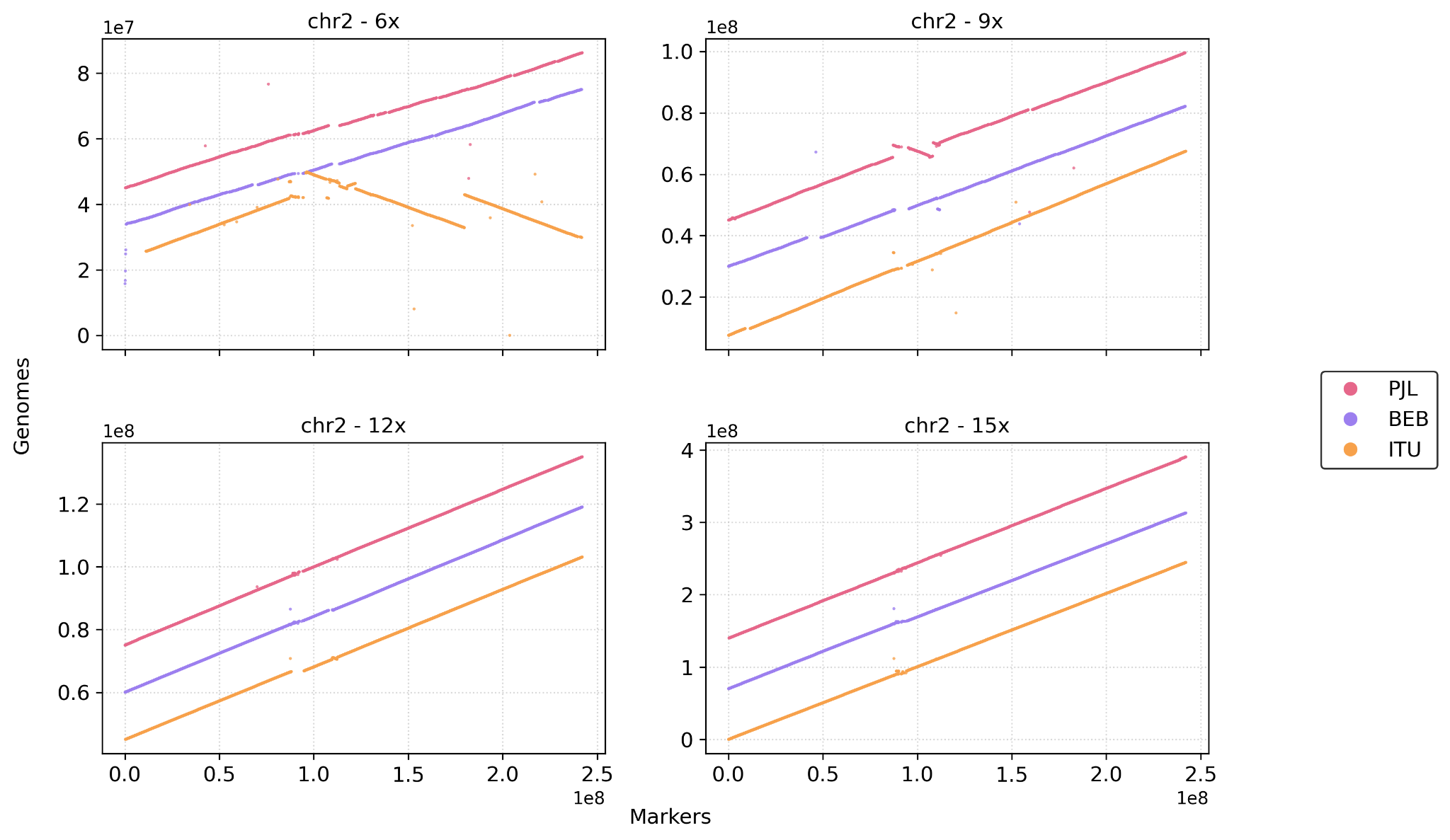

(B)

#chromosome 3

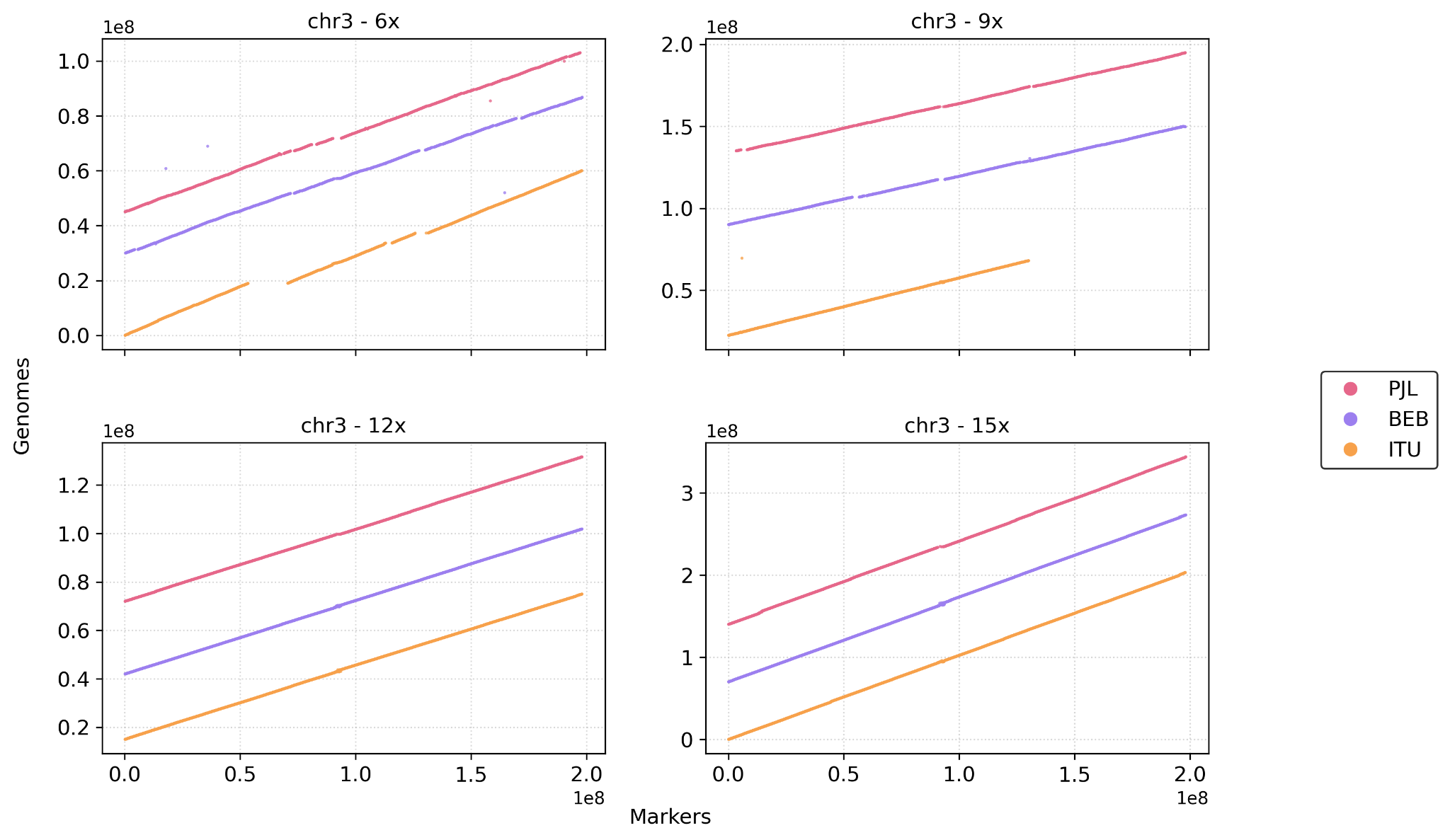

( C )

#chromosome 4

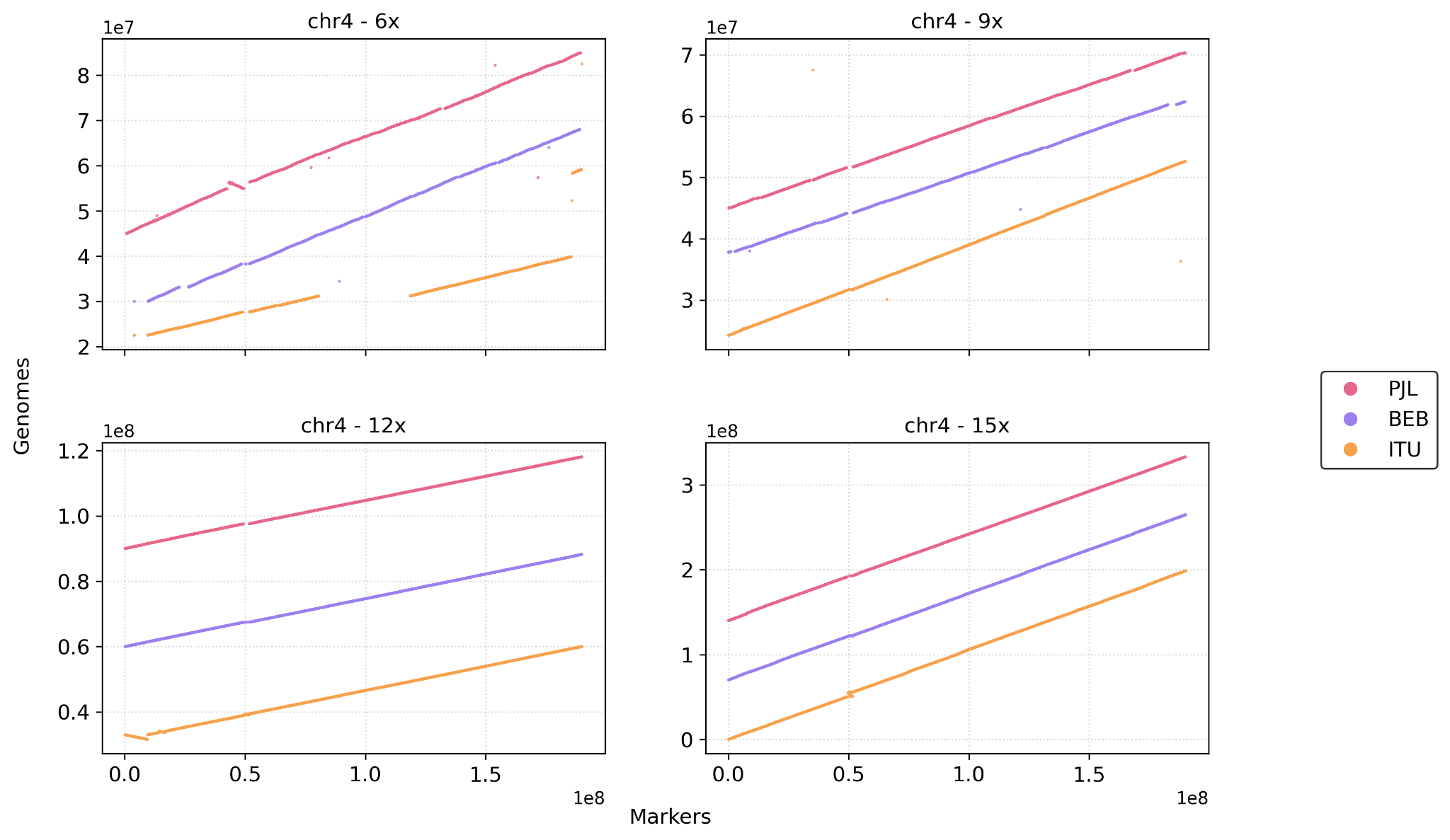

( D )

#chromosome 5

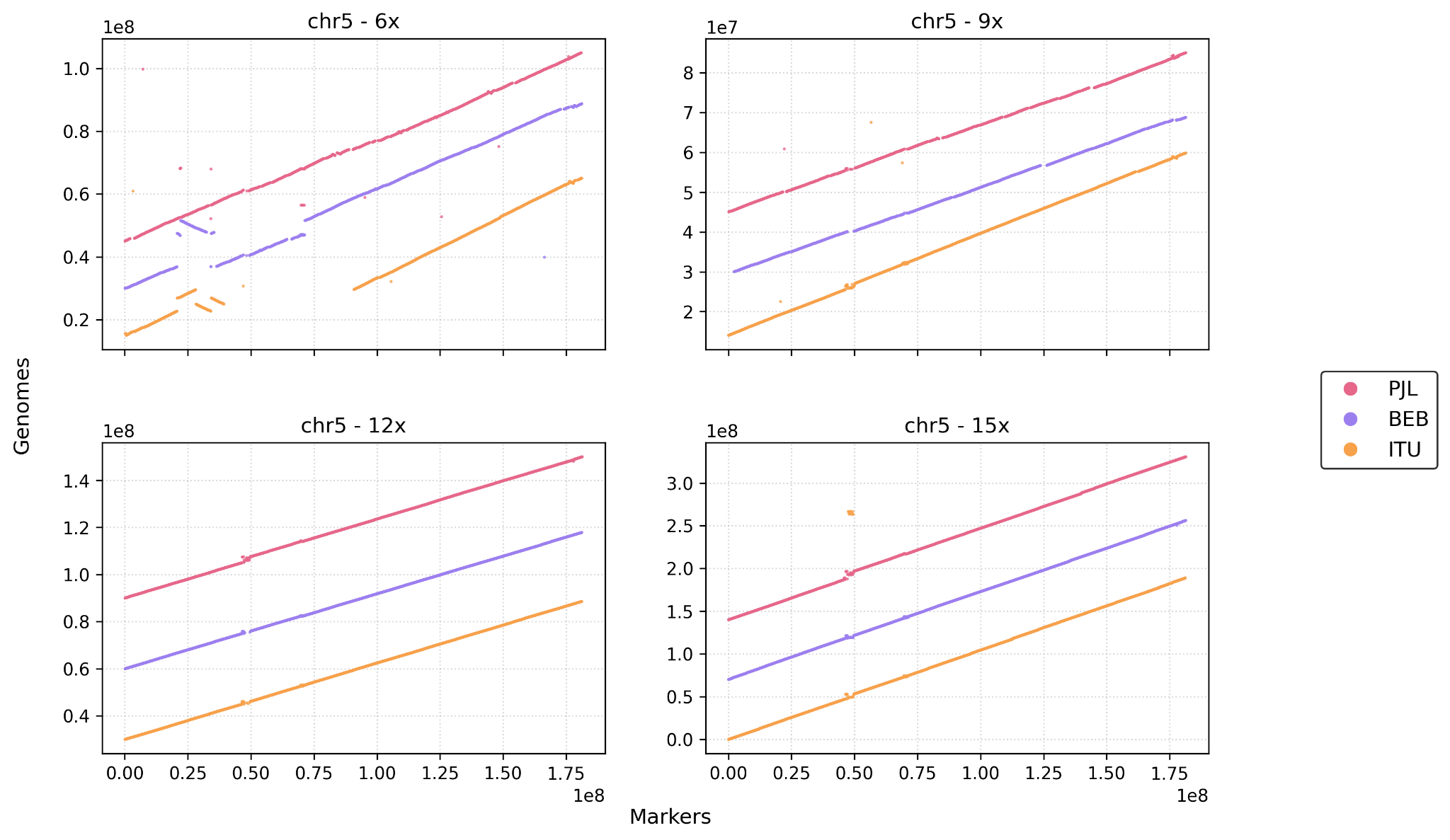

( E )

\# chromosome 6

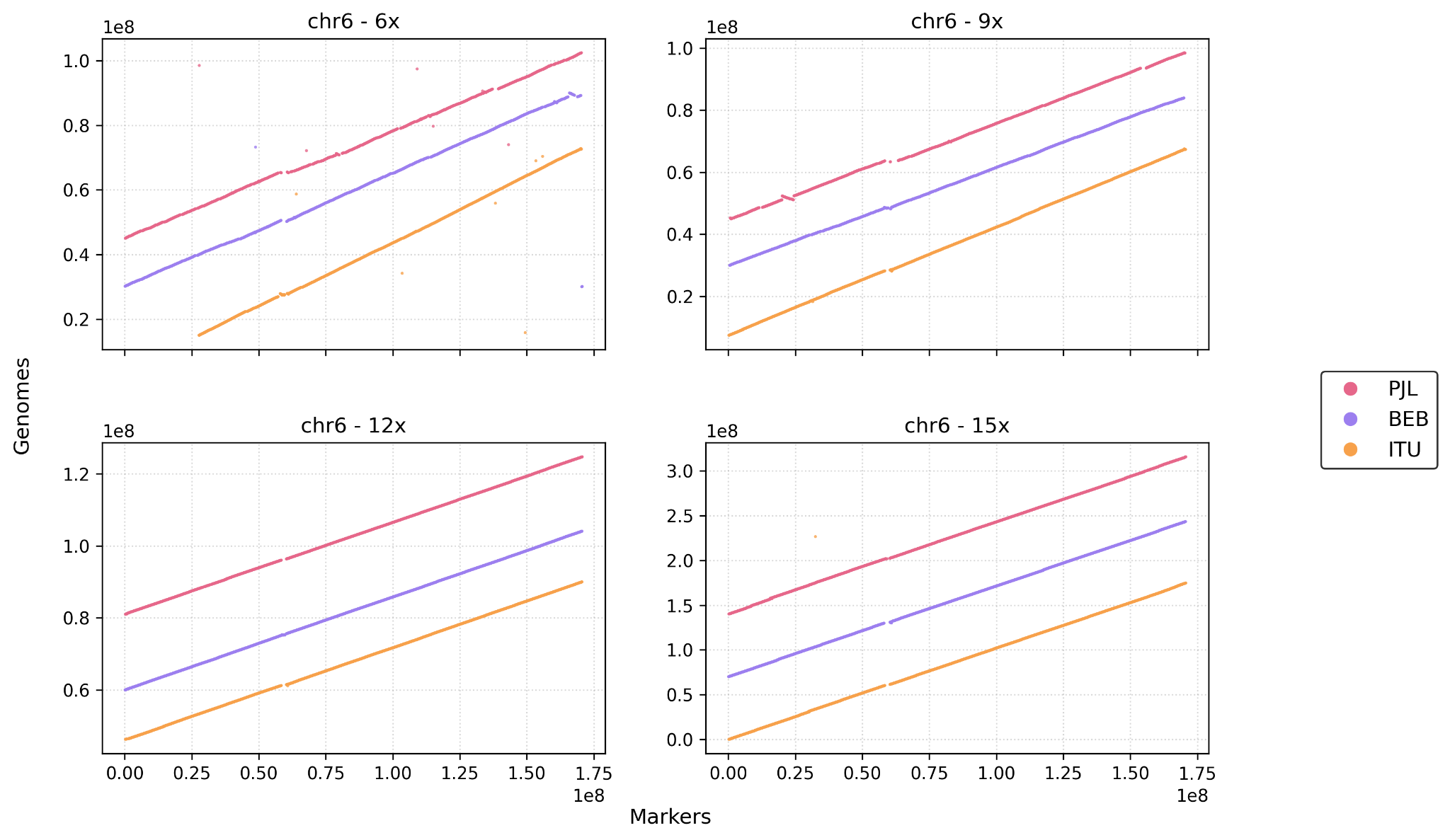

( F )

#chromosome 7

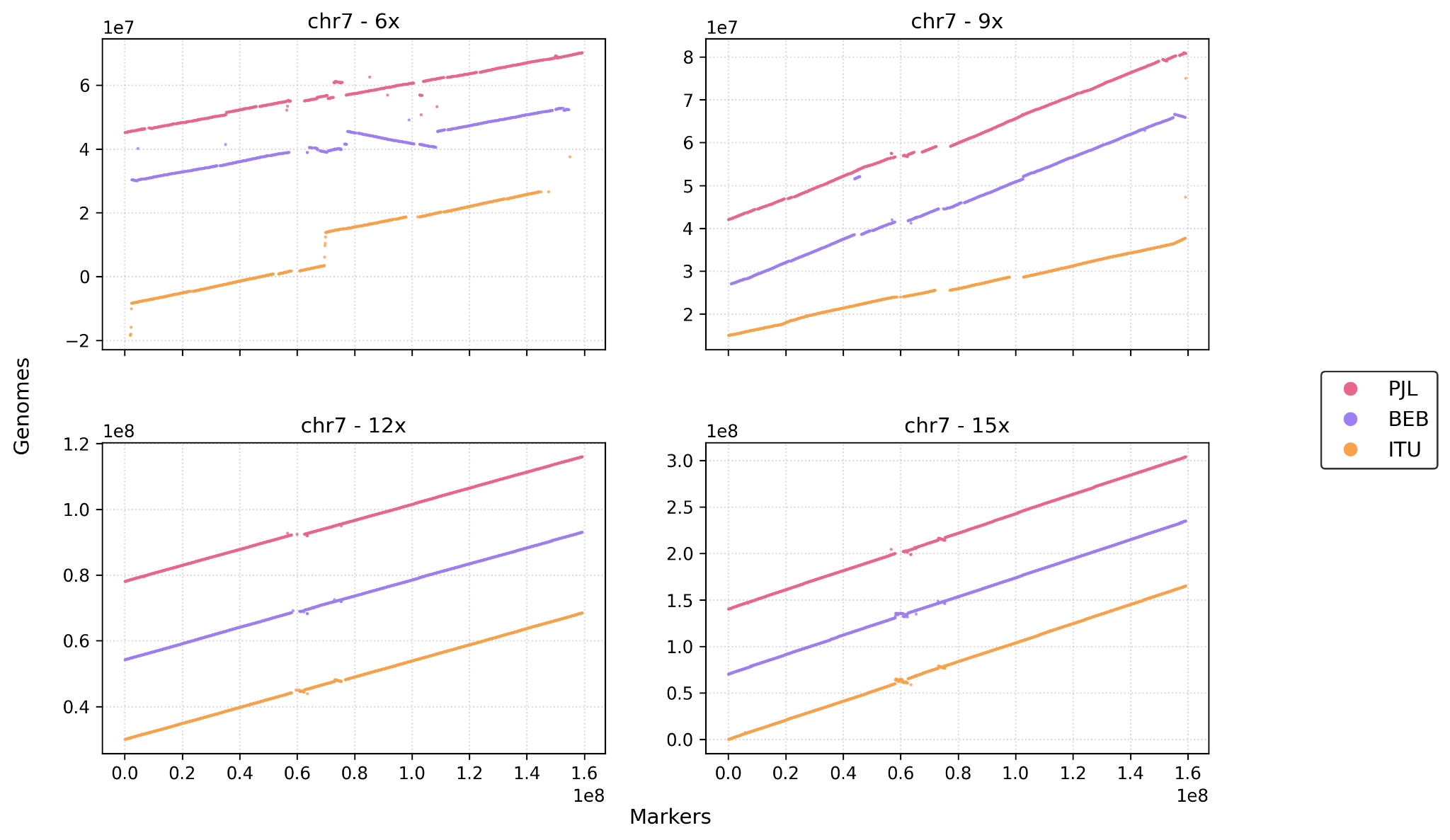

( G )

\# chromosome 8

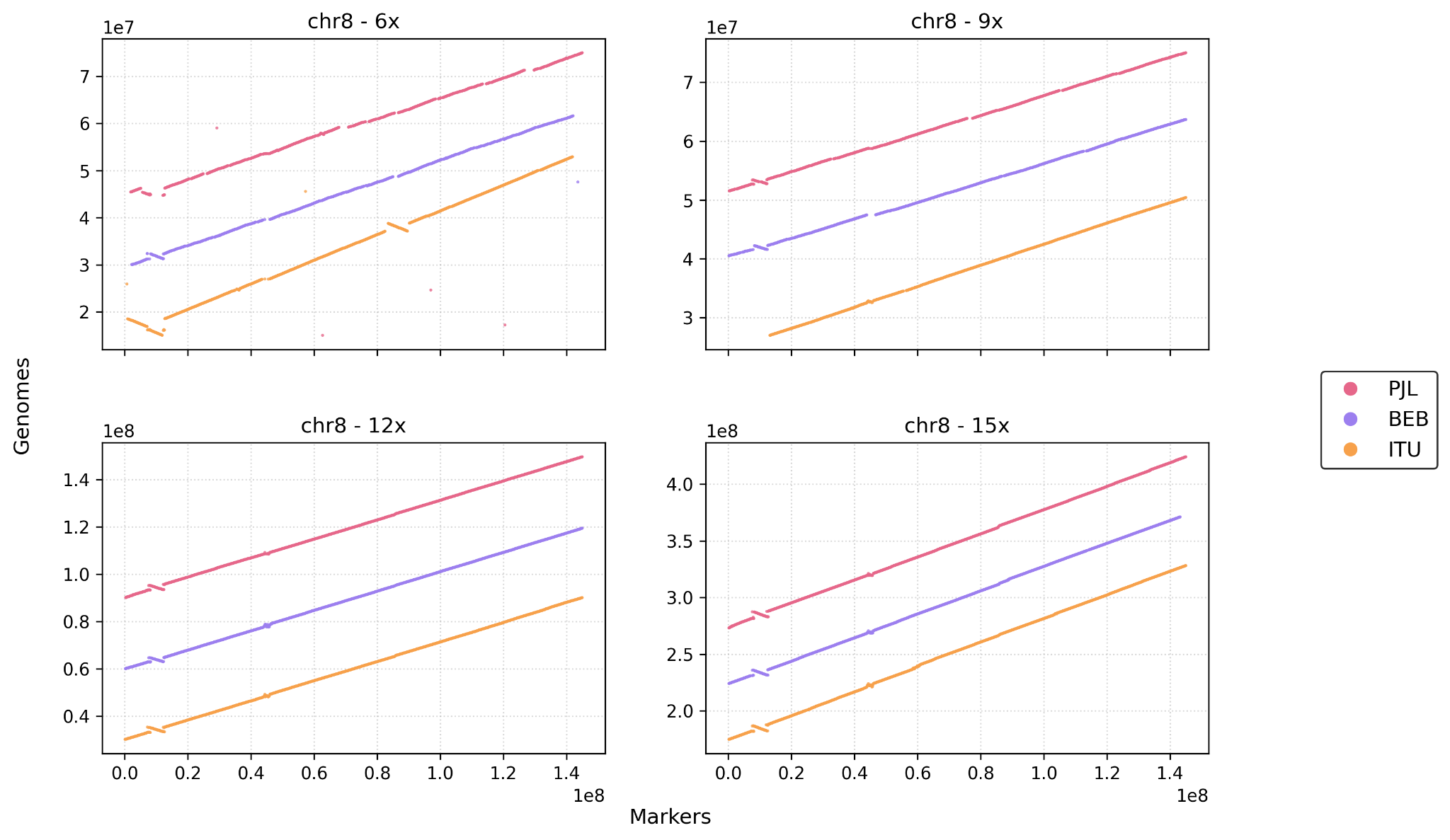

( H )

\# chromosome 9

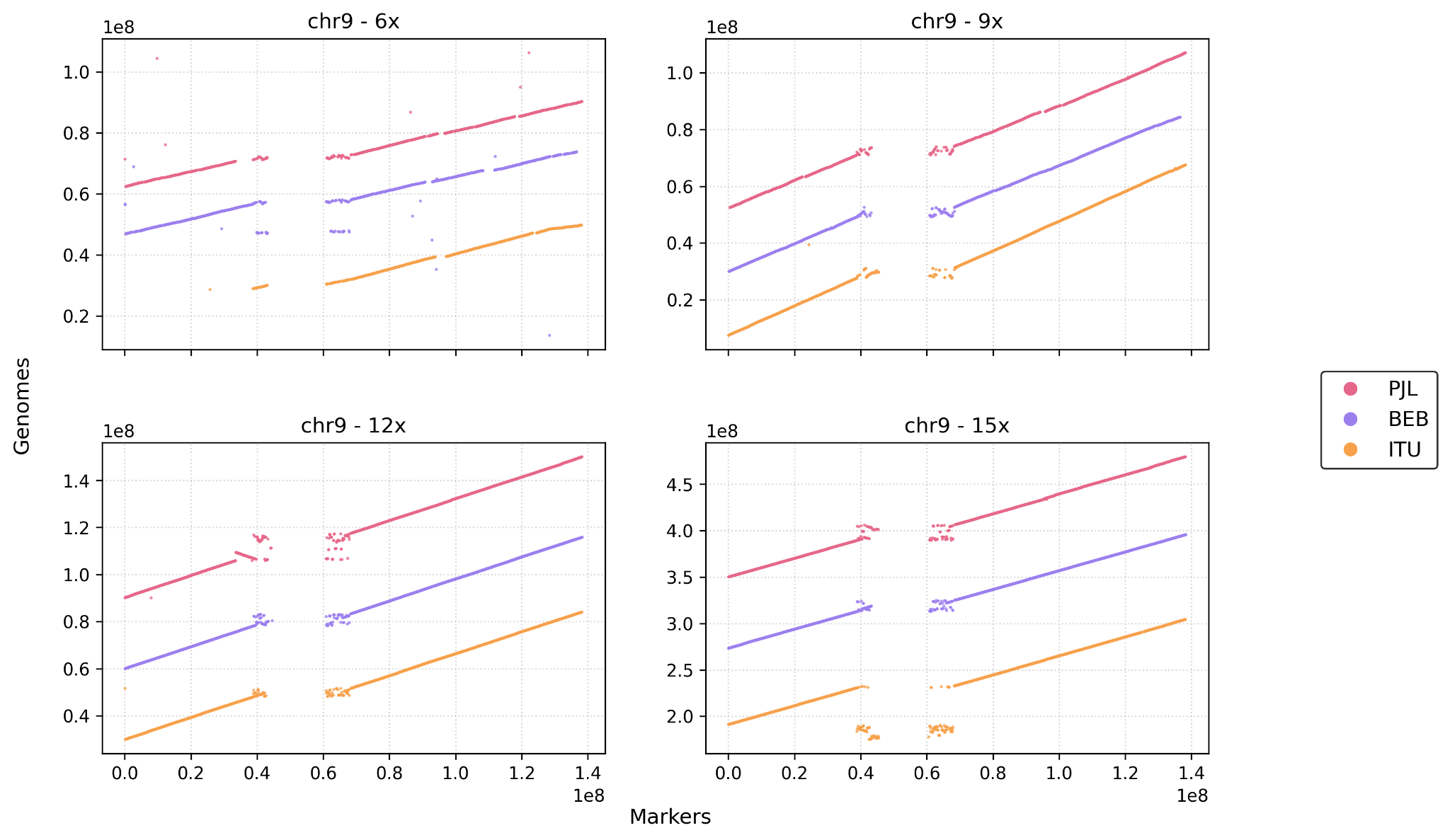

( I )

\# chromosome 10

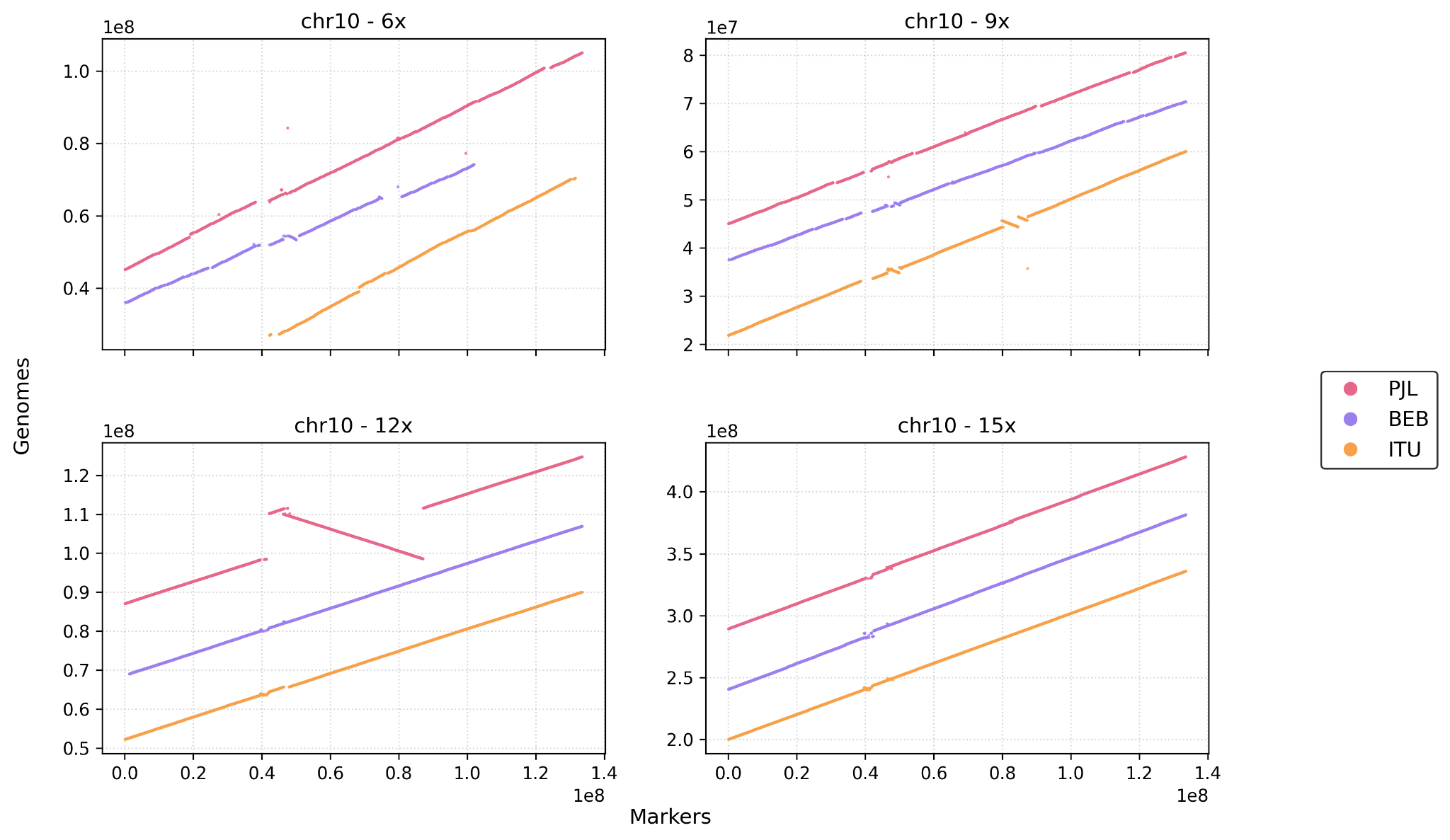

( J )

#chromosome 11

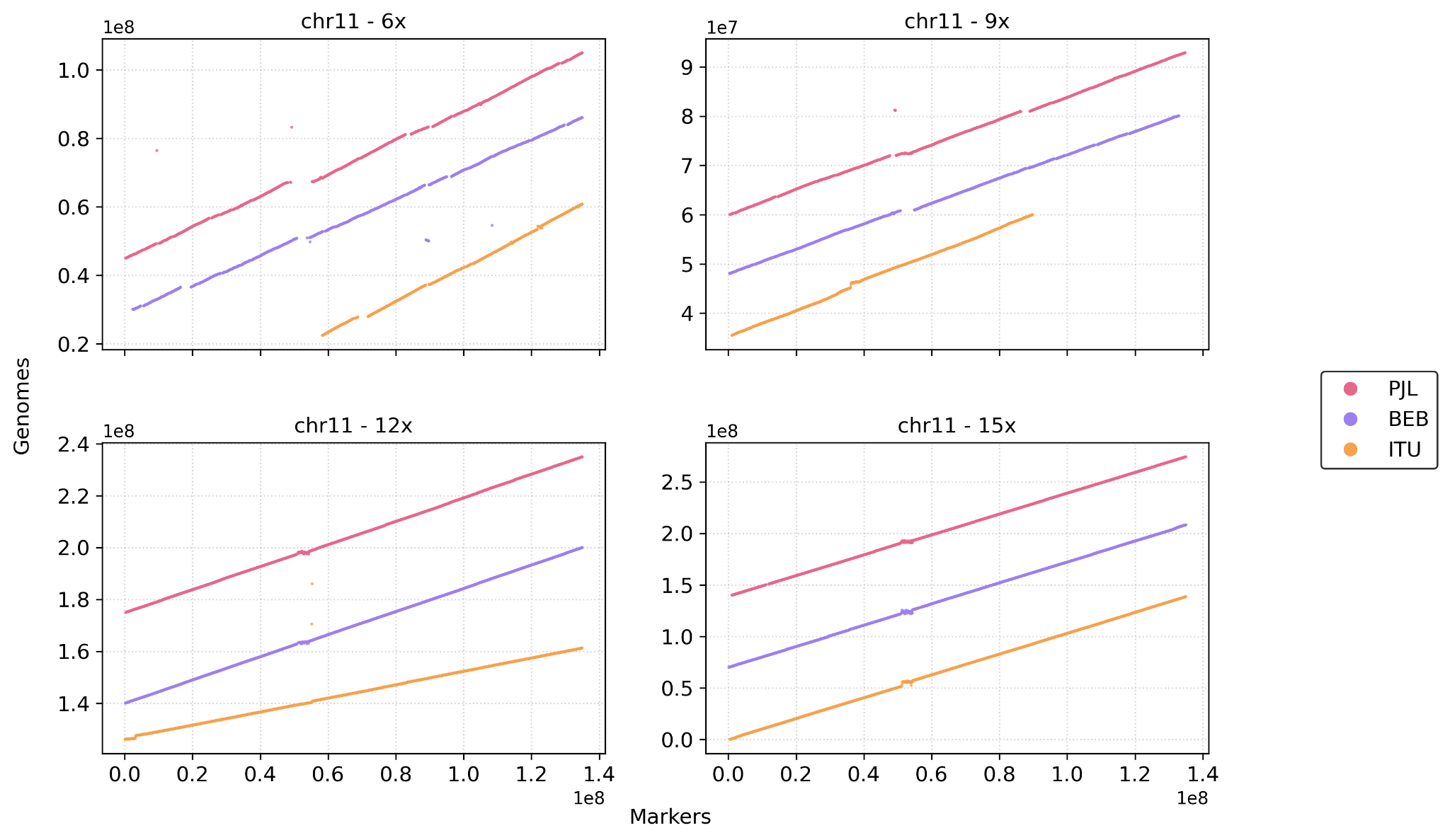

( K )

\# chromosome 12

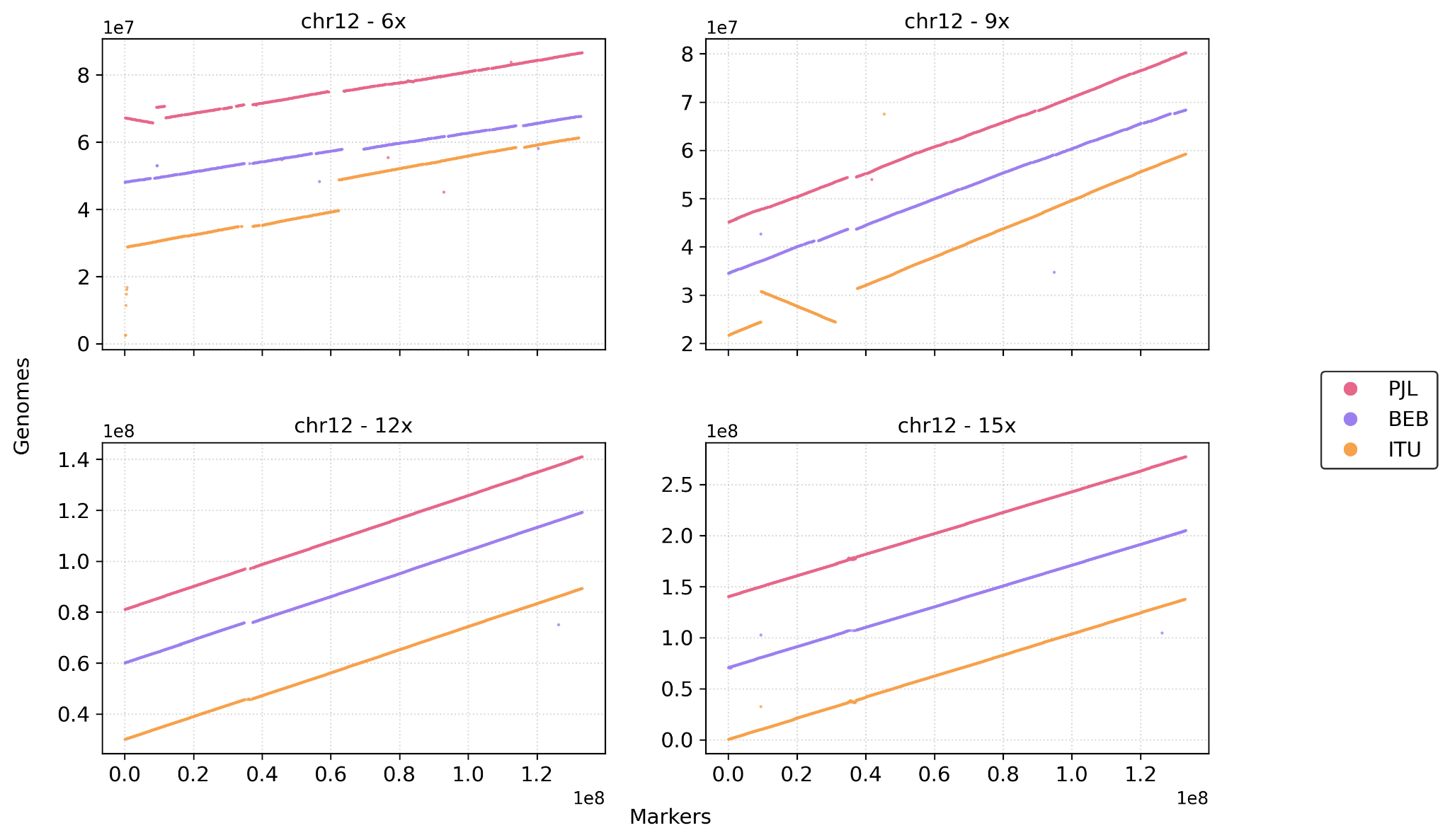

( L )

\# chromosome 13

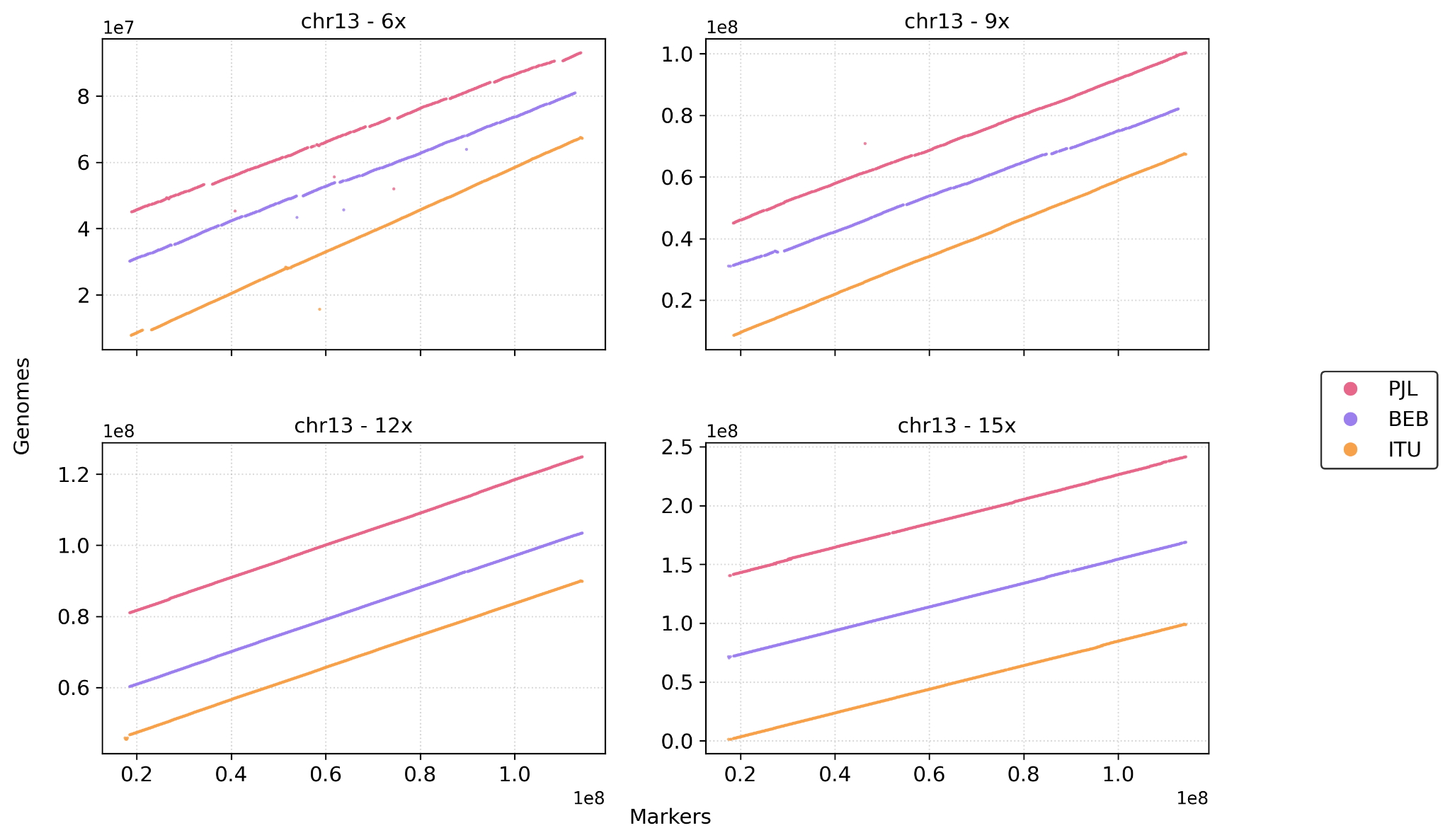

( M )

\# chromosome 14

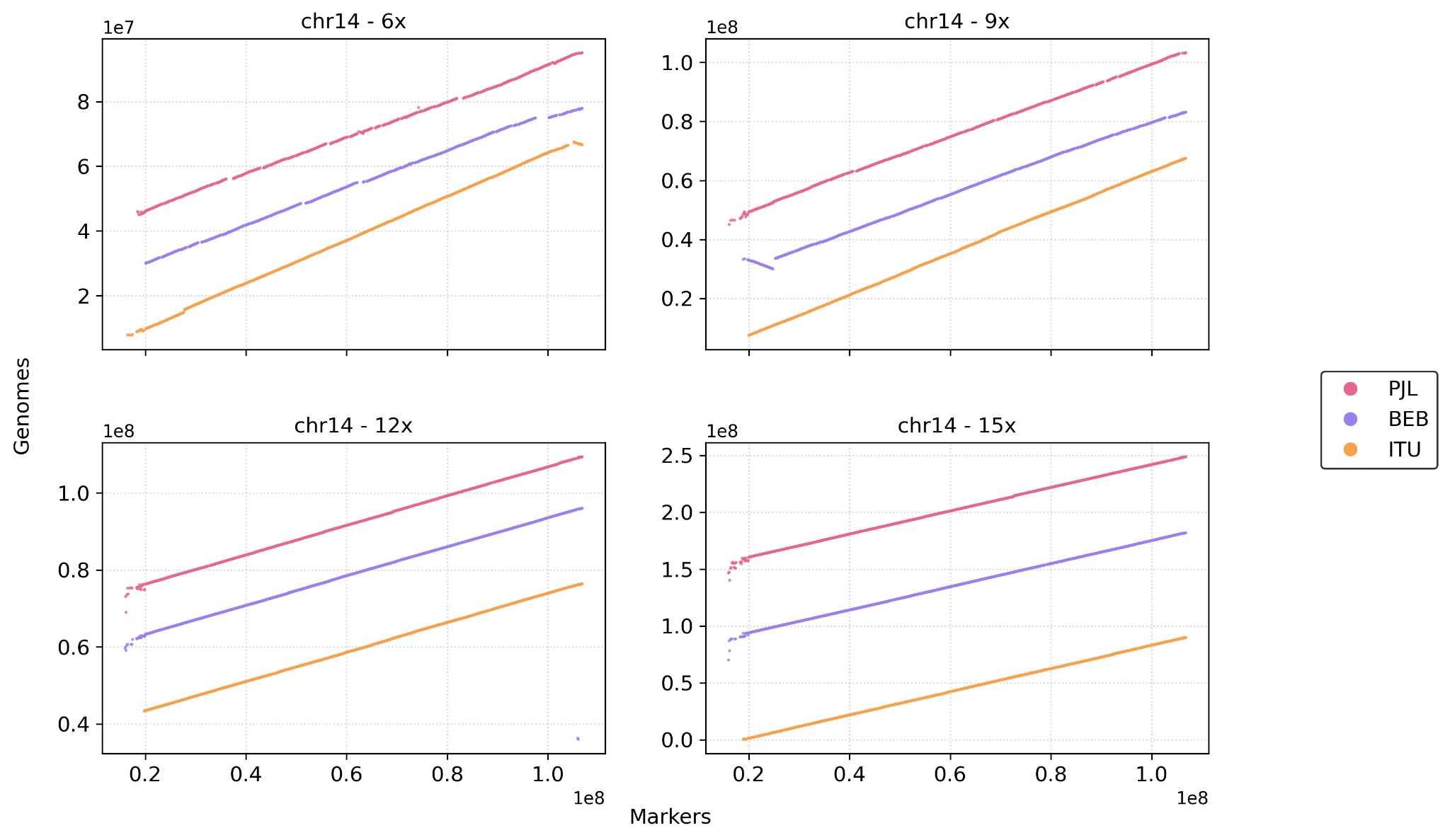

( N )

#chromosome 15

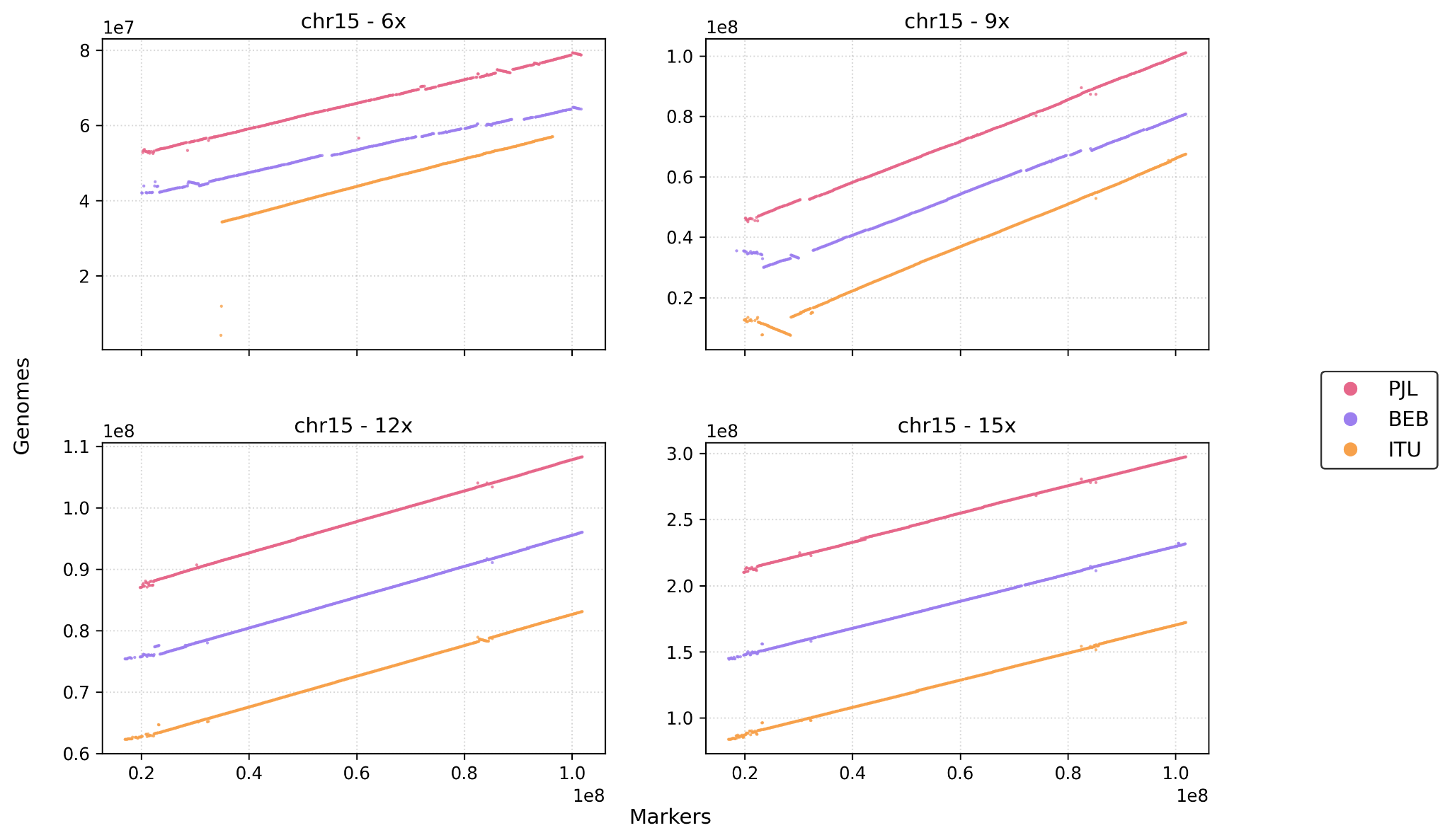

( O )

\# chromosome 16

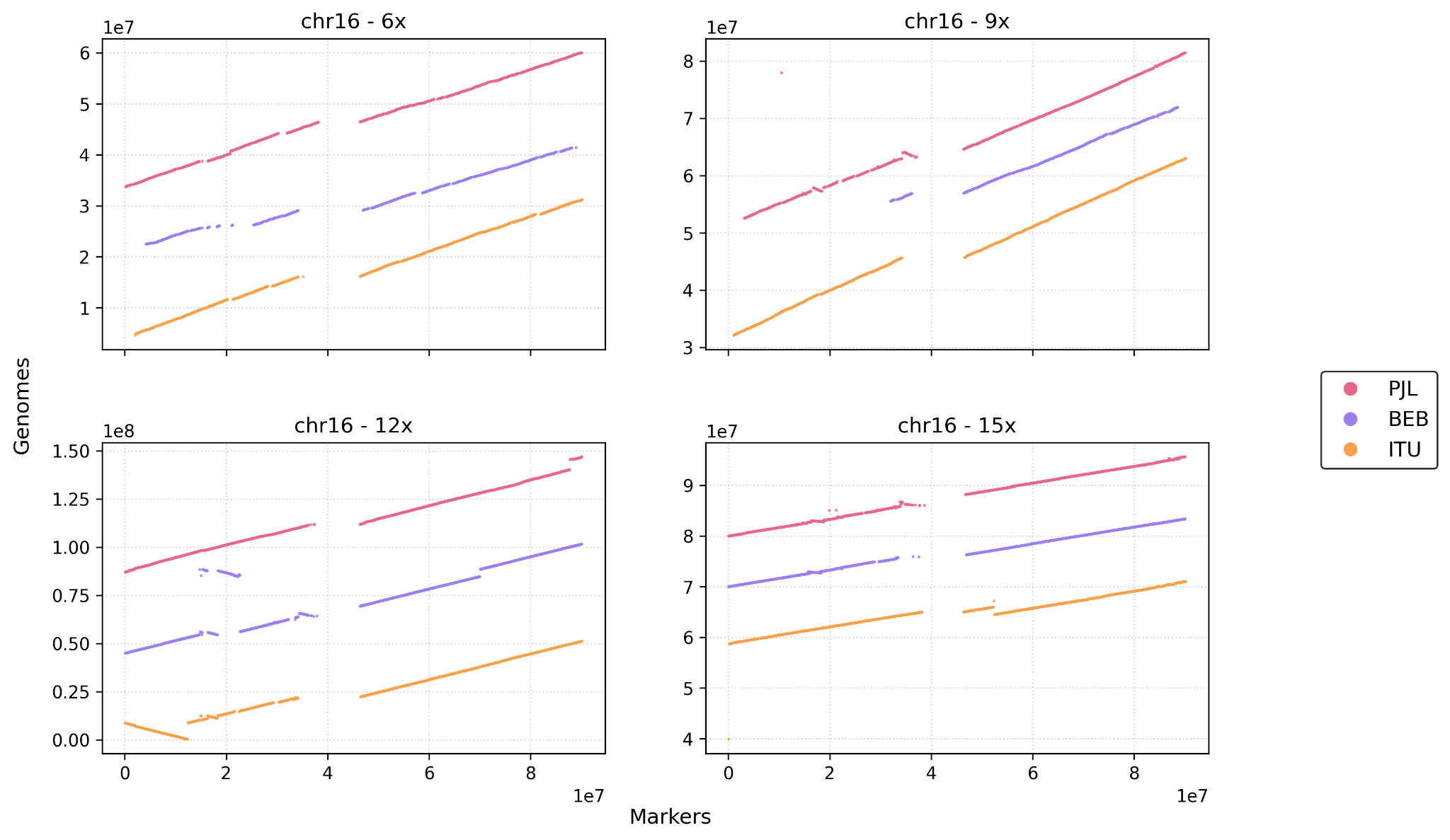

( P )

#chromosome 17

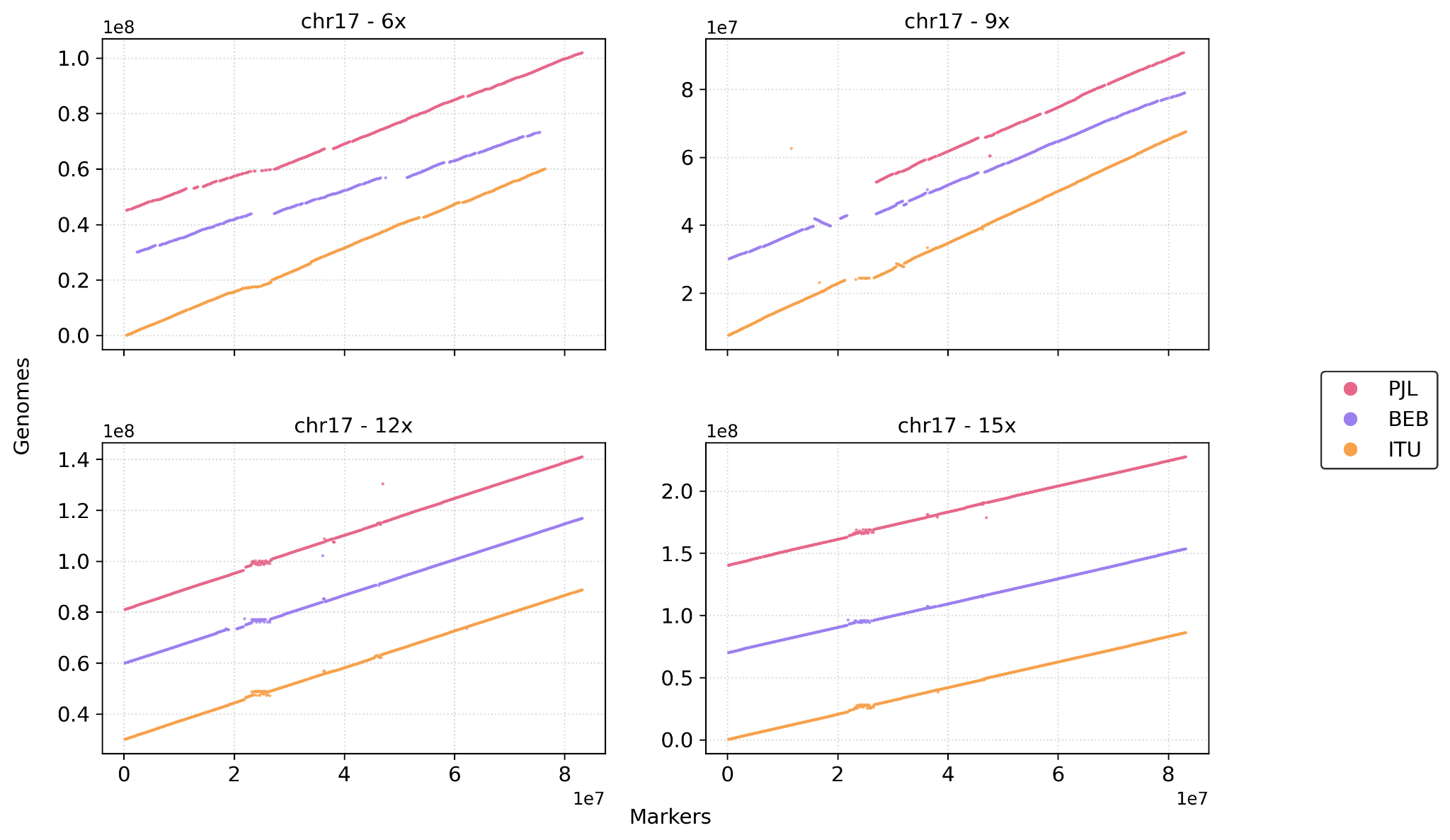

( Q )

\# chromosome 18

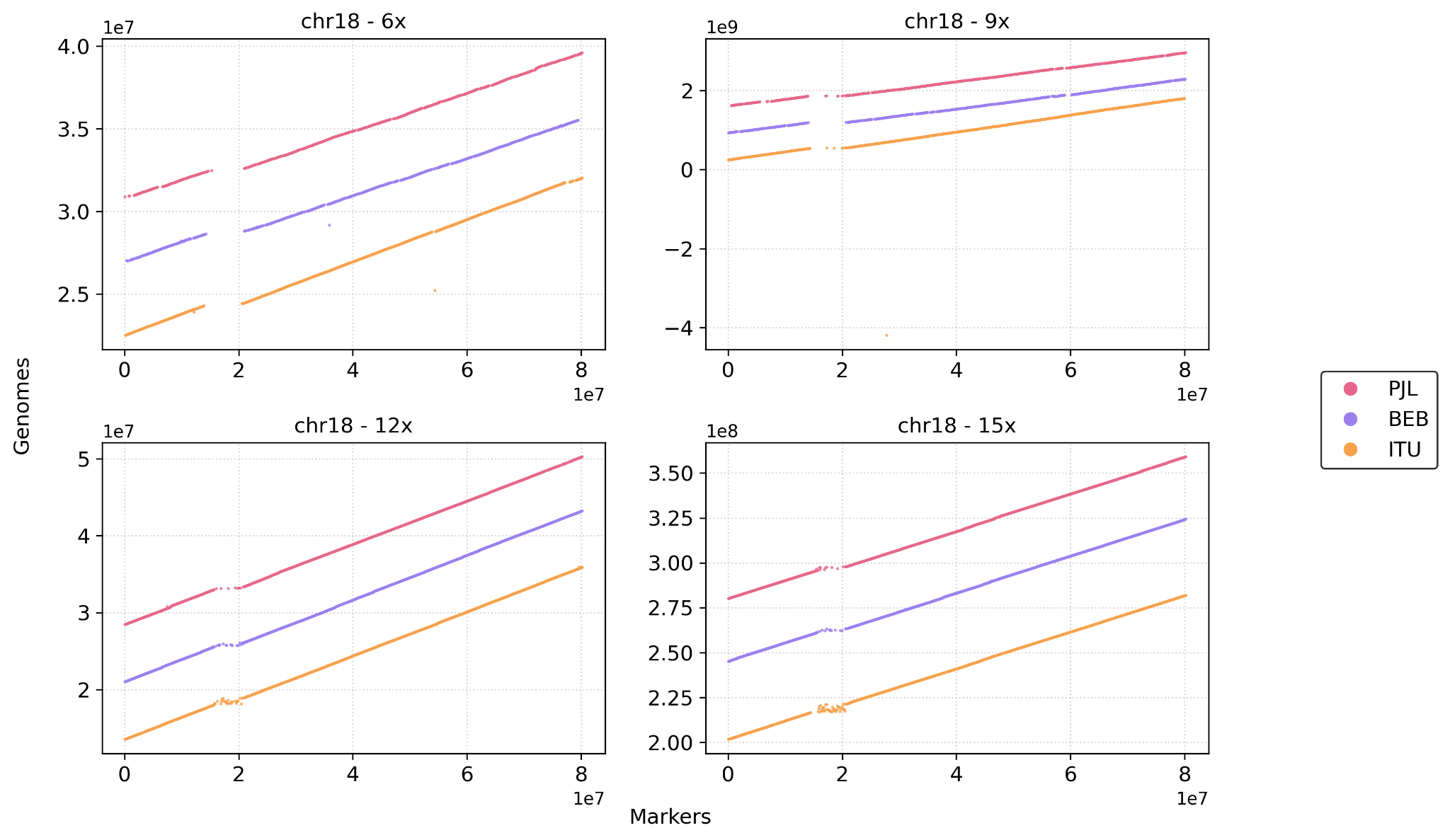

( R )

#chromosome 19

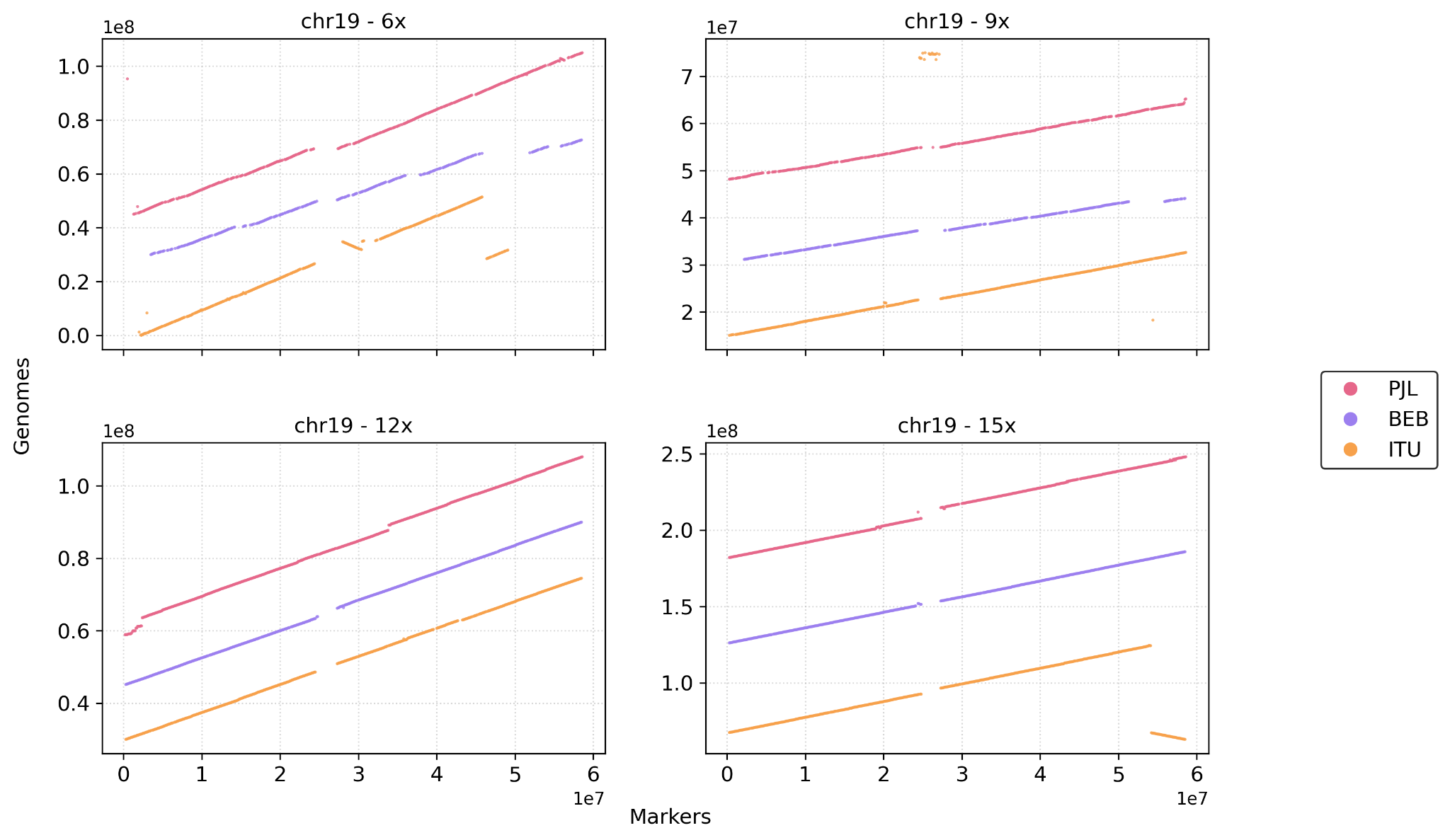

( S )

\# chromosome 20

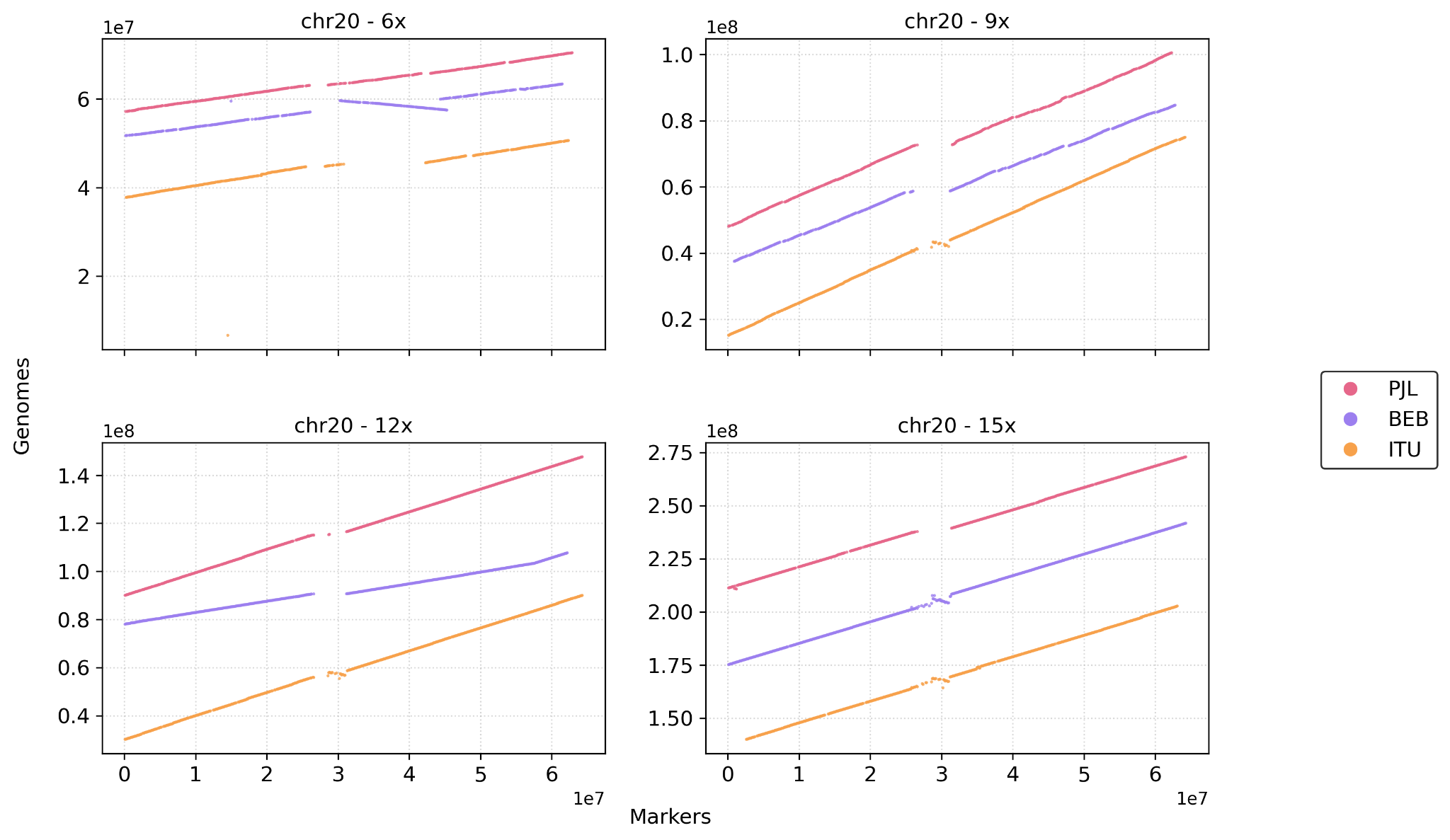

( T )

\# chromosome 21

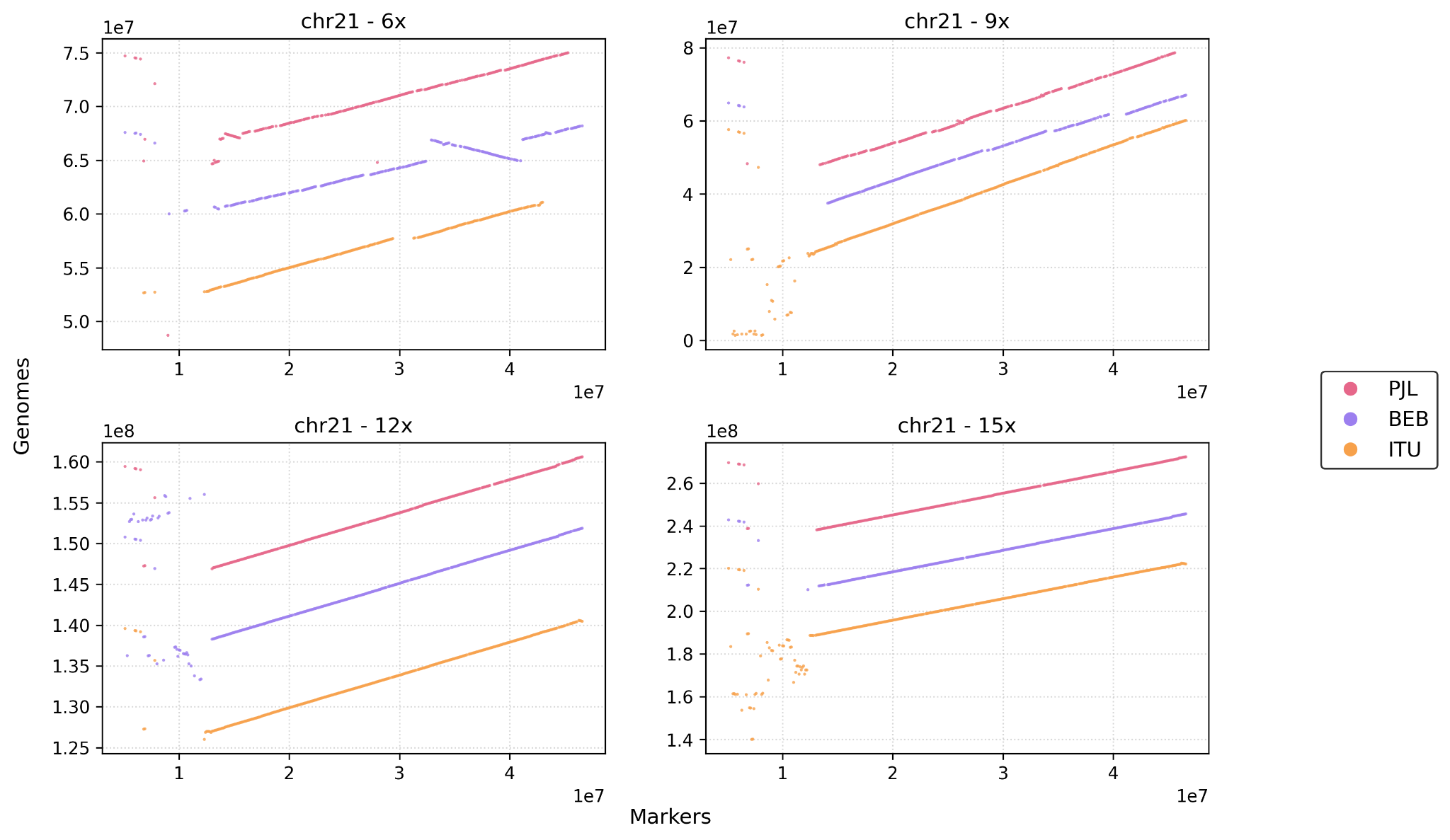

( U )

\# chromosome 22

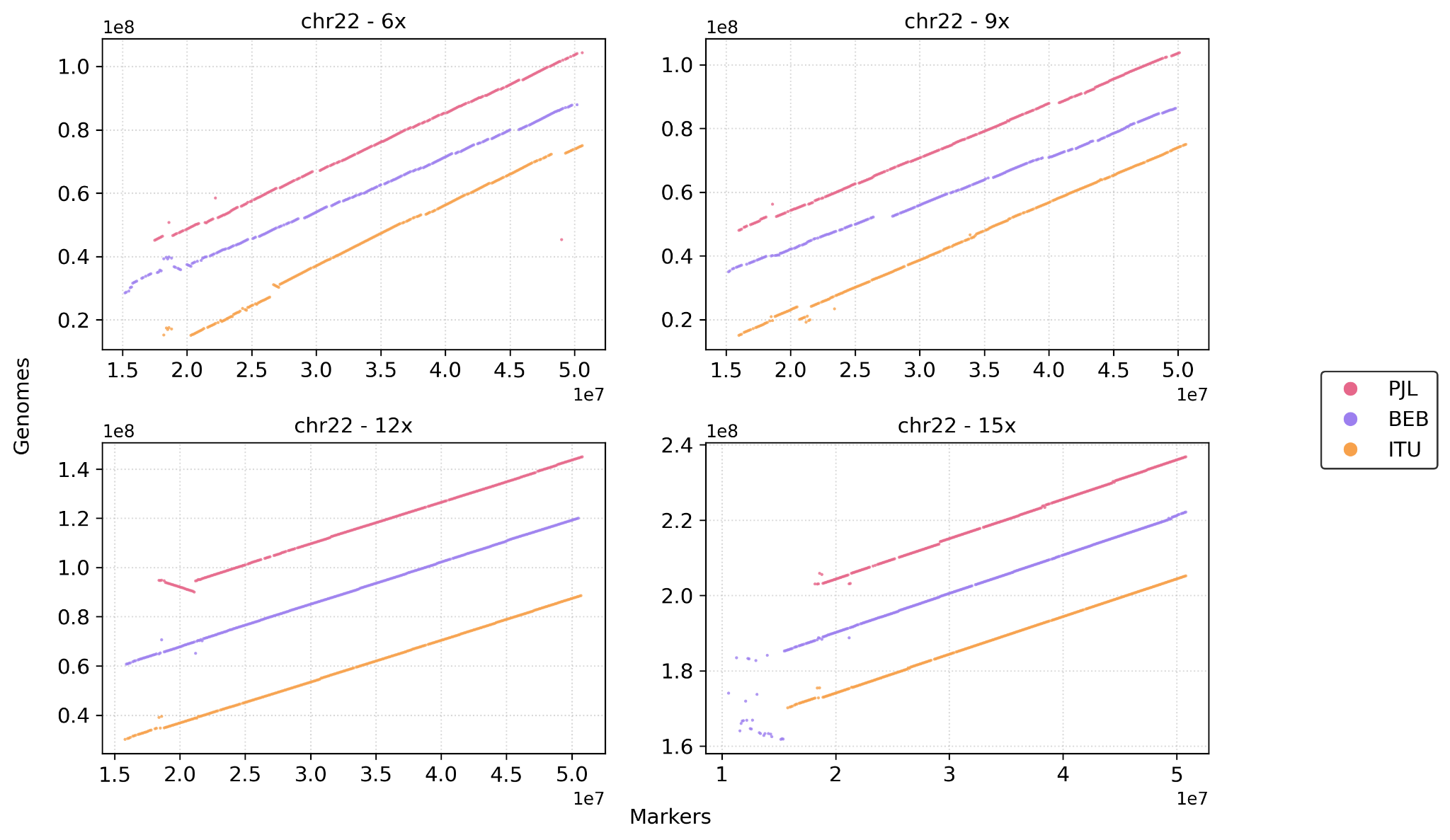

( V )

\# chromosome X

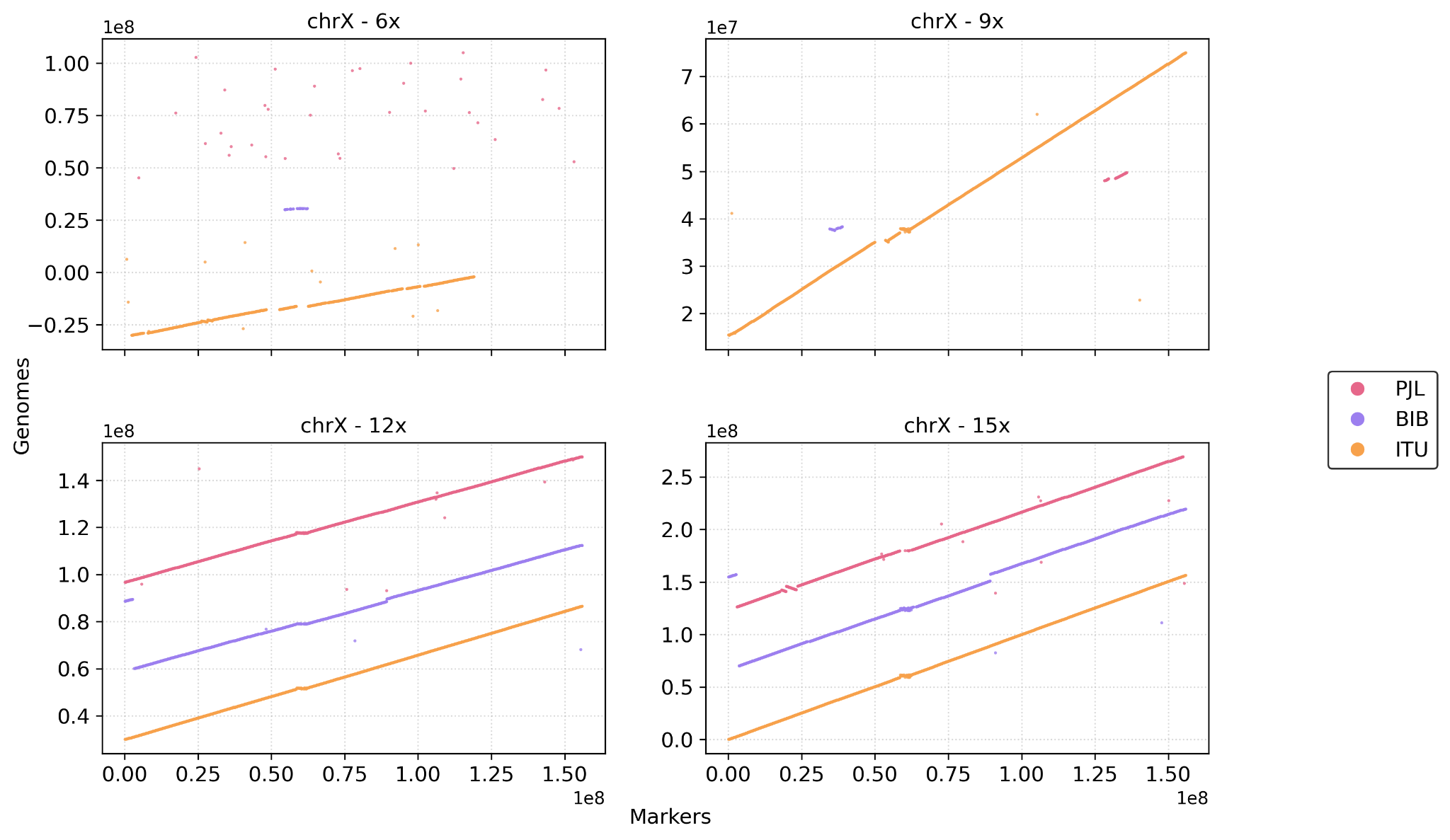

( W )

#### *Supplementary Figure S1 ( A - W )* : shows comparative dot plot of individual chromosomes from several genomes against virtual markers every 100kb from hg38 assembly.

**# Manual corrected chromosomes before correction**

#chr1_15x- ITU1

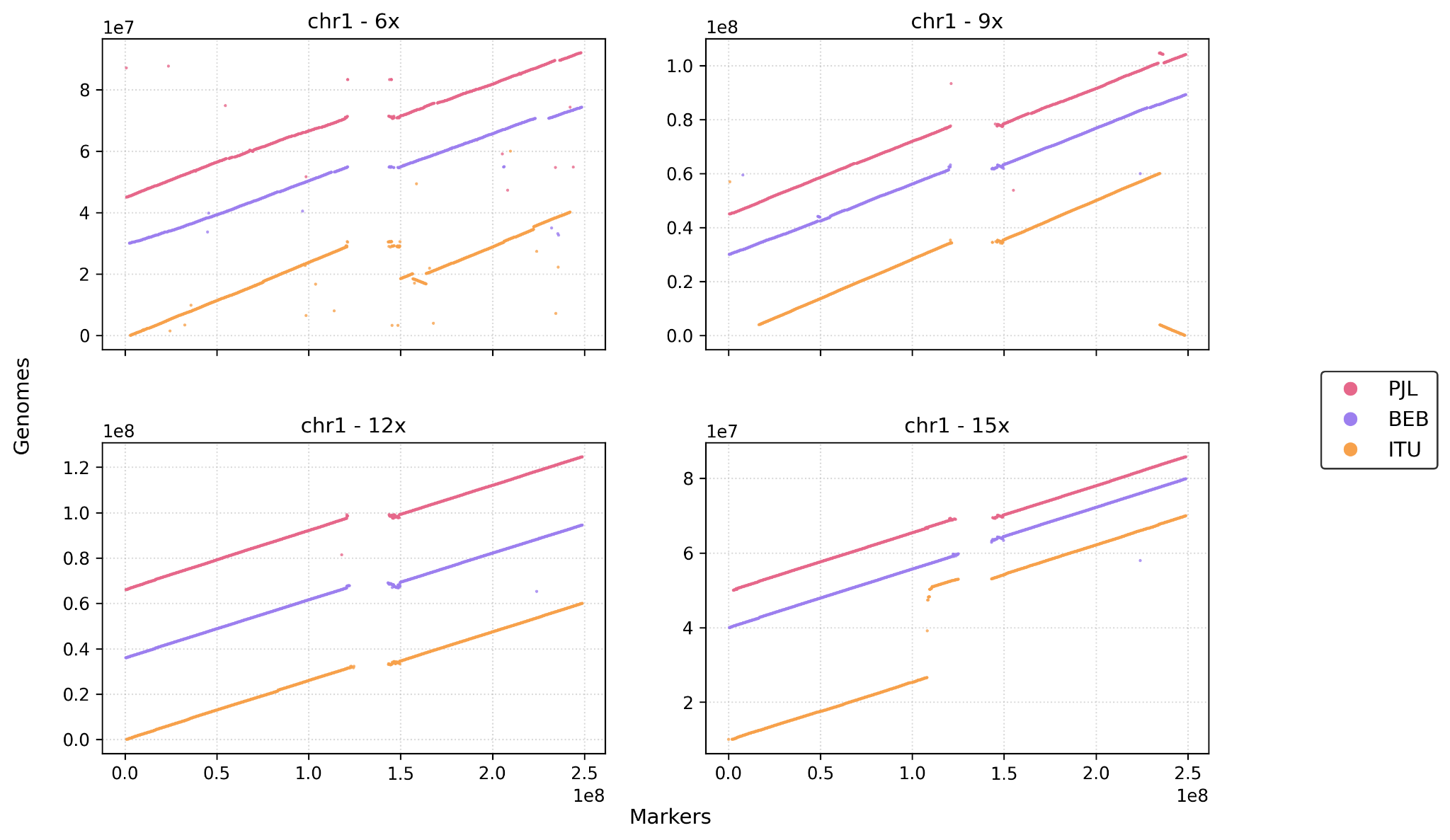

(i)

#chr19_12x- PJL1

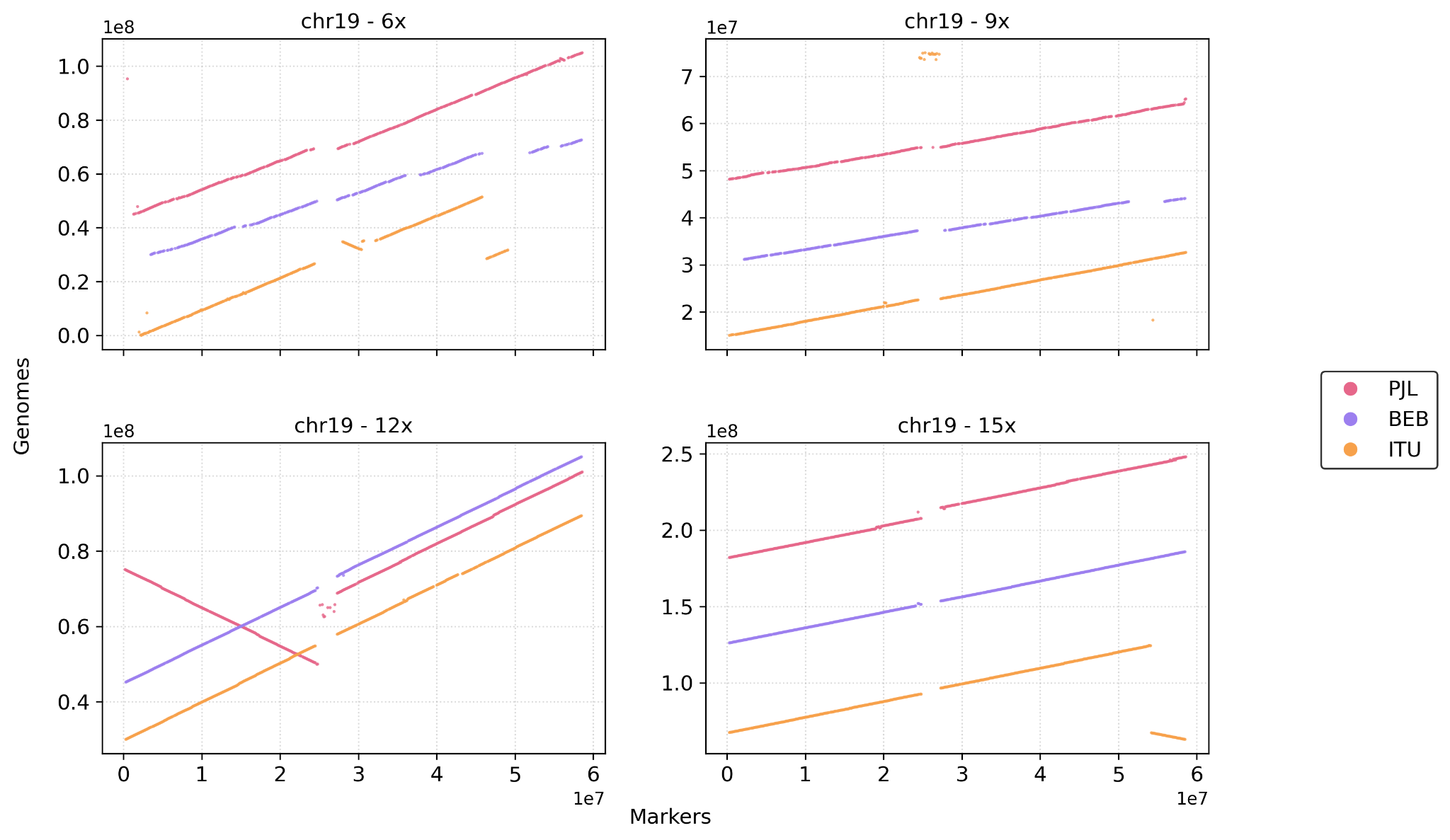

(ii)

#chr11_ITU-12x

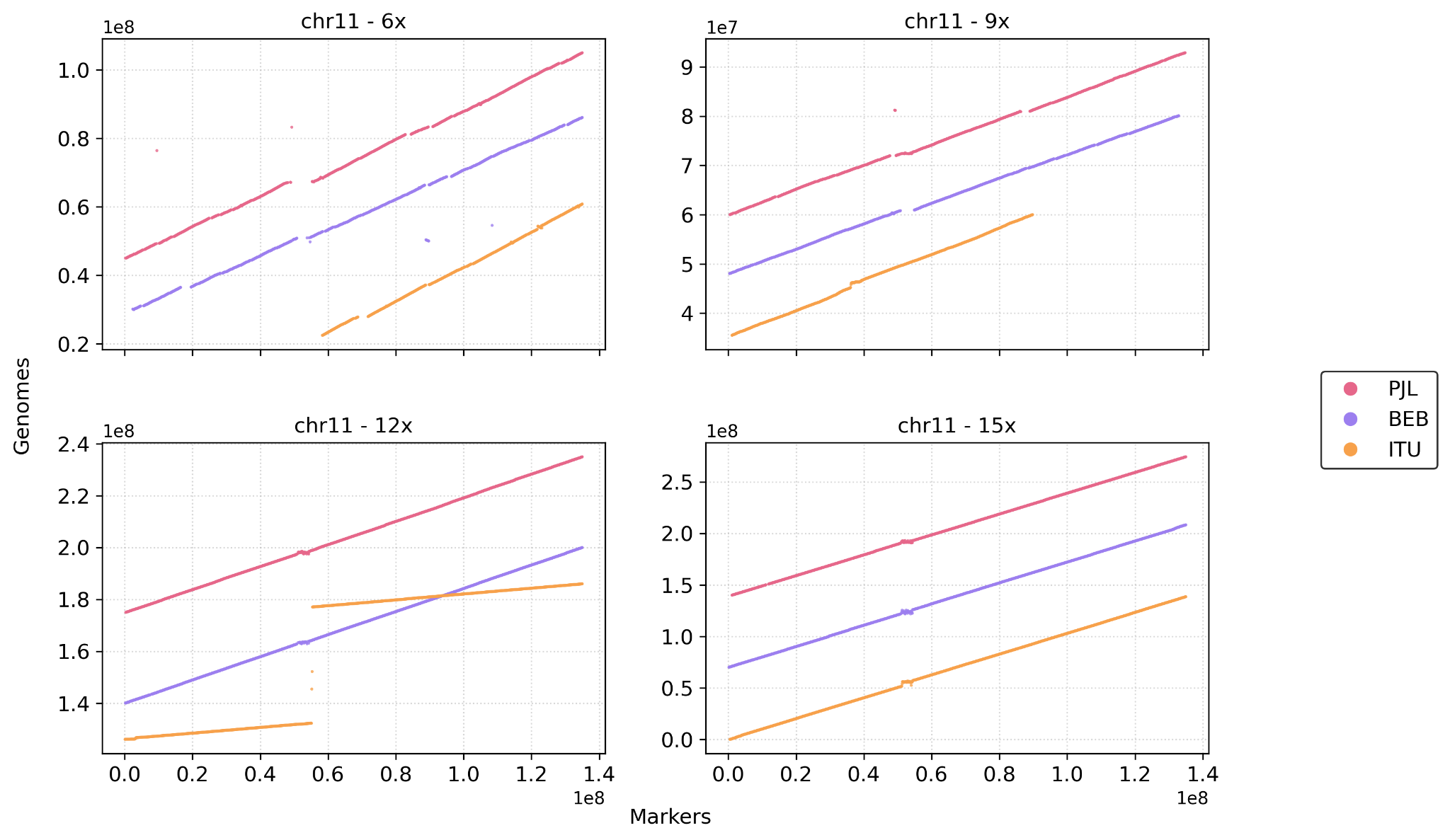

(iii)

#chr16_15x - ITU1

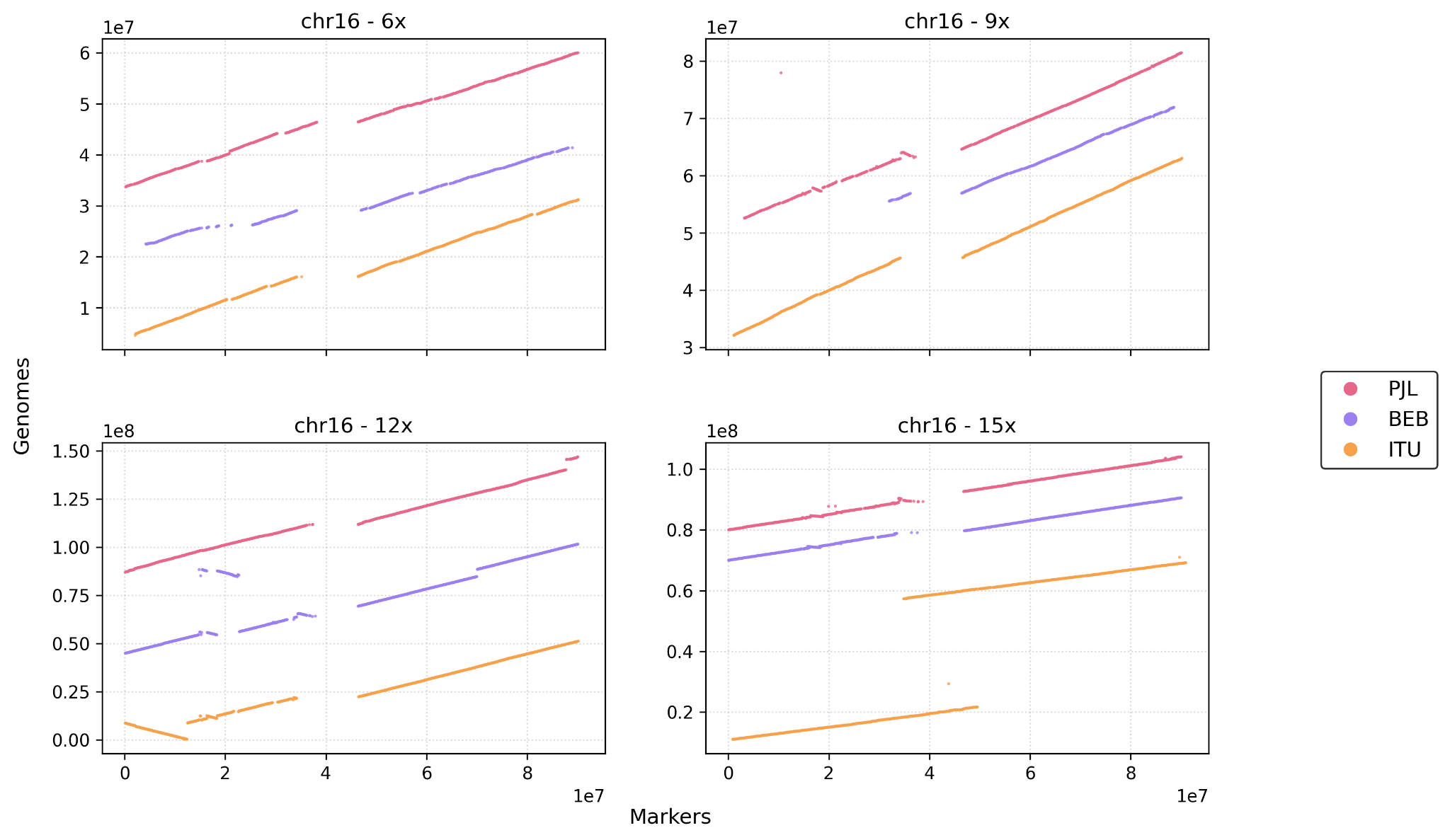

(iv)

#### *Supplementary Figure S2 ( i - iv ) : shows comparative dot plot of individual chromosomes from few genomes where p and q arms are splitting*
